# Amino acids 484 and 494 of SARS-CoV-2 spike are hotspots of immune evasion affecting antibody but not ACE2 binding

**DOI:** 10.1101/2021.04.22.441007

**Authors:** Marta Alenquer, Filipe Ferreira, Diana Lousa, Mariana Valério, Mónica Medina-Lopes, Marie-Louise Bergman, Juliana Gonçalves, Jocelyne Demengeot, Ricardo B. Leite, Jingtao Lilue, Zemin Ning, Carlos Penha-Gonçalves, Helena Soares, Cláudio M. Soares, Maria João Amorim

**Affiliations:** Cell Biology of Viral Infection Lab, Instituto Gulbenkian de Ciência, Oeiras, Portugal; ITQB NOVA, Instituto de Tecnologia Química e Biológica António Xavier, Universidade Nova de Lisboa; Oeiras, Portugal.; Lymphocyte Physiology Lab, Instituto Gulbenkian de Ciência; Oeiras, Portugal.; CEDOC NOVA, Centro de Estudos de Doenças Crónicas, Nova Medical School, Universidade Nova de Lisboa; Lisboa, Portugal.; Genomics Unit, Instituto Gulbenkian de Ciência; Oeiras, Portugal.; Bioinformatics Unit, Instituto Gulbenkian de Ciência; Oeiras, Portugal.; The Wellcome Trust Sanger Institute; Hinxton, UK.; Disease Genetics Lab, Instituto Gulbenkian de Ciência; Oeiras, Portugal.

**Author notes:** These authors contributed equally to this work (Filipe Ferreira, Diana Lousa).

## Abstract

Understanding SARS-CoV-2 evolution and host immunity is critical to control COVID-19 pandemics. At the core is an arms-race between SARS-CoV-2 antibody and angiotensin-converting enzyme 2 (ACE2) recognition, a function of the viral protein spike. Mutations in spike impacting antibody and/or ACE2 binding are appearing worldwide, with the effect of mutation synergy still incompletely understood. We engineered 25 spike-pseudotyped lentiviruses containing individual and combined mutations, and confirmed that E484K evades antibody neutralization elicited by infection or vaccination, a capacity augmented when complemented by K417N and N501Y mutations. *In silico* analysis provided an explanation for E484K immune evasion. E484 frequently engages in interactions with antibodies but not with ACE2. Importantly, we identified a novel amino acid of concern, S494, which shares a similar pattern. Using the already circulating mutation S494P, we found that it reduces antibody neutralization of convalescent and post-immunization sera, particularly when combined with E484K and N501Y. Our analysis of synergic mutations provides a landscape for hotspots for immune evasion and for targets for therapies, vaccines and diagnostics.

**One-Sentence Summary:** Amino acids in SARS-CoV-2 spike protein implicated in immune evasion are biased for binding to neutralizing antibodies but dispensable for binding the host receptor angiotensin-converting enzyme

## Main Text

Severe acute respiratory syndrome coronavirus–2 (SARS-CoV-2) is the virus responsible for the pandemic of coronavirus disease 2019 (COVID-19) ^1^ that has caused more than 140 million infections and provoked the death of over 3 million people (as of April 18, 2021). A pertinent question is whether viral evolution will permit escaping immunity developed by natural infection or by vaccination. The answer to this multi-layered and complex question depends on the type, duration and heterogeneity of the selective pressures imposed by the host and by the environment to the virus, but also on the rate and phenotypic impact of the mutations the virus acquires. The central question is whether the virus is able to select mutations that escape host immunity while remaining efficient to replicate in the host ^2–4^. Identifying hotspots of viral immune evasion is critical for preparedness of future interventions to control COVID-19.

SARS-CoV-2 is an enveloped virus, characterized by displaying spike proteins at the surface ^5^. Spike is critical for viral entry ^6^ and is the primary target of vaccines and therapeutic strategies, as this protein is the immunodominant target for antibodies ^7–11^. Spike is composed of S1 and S2 subdomains. S1 contains the N-terminal (NTD) and receptor-binding (RBD) domains, and the S2 contains the fusion peptide (FP), heptad repeat 1 (HR1) and HR2, the transmembrane (TM) and cytoplasmic domains (CD) ^12^. S1 leads to the recognition of the angiotensin-converting enzyme 2 (ACE2) receptor and S2 is involved in membrane fusion ^6, 13, 14^. Spike binds to ACE2 displayed at the host cell surface, followed by proteolytic cleavage to yield S1 and S2 fragments ^6^. Interestingly, the spike protein oscillates between distinct conformations to recognize and bind different host factors ^15, 16^. In the prefusion state, the RBD domain alternates between open (‘up’) and closed (‘down’) conformations ^5, 17^. For binding to the ACE2 receptor, the RBD is transiently exposed in the ‘up’ conformation. However, most potent neutralizing antibodies elicited by natural infection ^9, 11, 18^ and by vaccination ^7, 10^ bind spike in the closed (‘down’) conformation. Understanding how each amino acid of spike dynamically interacts with either antibodies or ACE2 receptor may reveal the amino acid residues that are more prone to suffer mutations driving immunological escape without affecting viral entry.

Viral mutations may affect host-pathogen interactions in many ways: affect viral spread, impact virulence, escape natural or vaccine-induced immunity, evade therapies or detection by diagnostic tests, and change host species range ^19, 20^. Therefore, it is critical to survey circulating variants and assess their impact in the progression of SARS-CoV-2 dynamics in the population, in real time. Being a novel virus circulating in the human population, SARS-CoV-2 evolution displayed a mutational pattern of mostly neutral random genetic drift until December 2020. In fact, the D614G mutation in the spike protein was amongst the few epidemiologically significant variants, resulting in increased transmissibility without affecting the severity of the disease ^21, 22^. However, from the end of 2020, three divergent SARS-CoV-2 lineages evolved into fast-spreading variants that became known as variants of concern B.1.1.7 [United Kingdom (UK)] ^23^, B.1.351 [South Africa (SA)] ^24^ and P.1 (Brazil) ^25^. More recently, other lineages were added to this list: B.1.427, B.1.429 [California, United States of America (USA)], B.1.617 or B.1.617.1-3 [India] ^23, 26, 27^, as well as several lineages of interest that carry the mutation E484K, including P.2 ^28^ and B.1.1.7 containing E484K. These variants present alterations in the spike protein that change its properties: increase transmissibility and virulence ^29, 30^ and/or escape immunity developed by natural infection ^31, 32^, therapies ^33–35^, vaccination ^36^, detection ^37^; and change the host species range ^38^. Mutations in the RBD of spike may severely affect viral replication and host immune response since, as explained above, this region is responsible for binding to ACE2 and is immunodominant. Three mutations D614G, N501Y and L452R are associated with an increased ACE2 binding affinity in humans and increased viral transmission ^21, 22, 29, 39^. Y453F was associated with a mink-to-human adaptation in cluster 5 ^40^. L452R and N439K are associated with a modest reduction in antibody-dependent neutralization by immune sera, whilst variants containing E484K display a reduction that is moderate to substantial ^41^. Variants, however, contain several other mutations. Interestingly, some mutations are convergent whilst others are unique in lineages ^42^. For example, in B.1.351 and P.1 lineages, mutations in amino acid residues at positions 18 (L18F) and at position 417 (S417N, or S414T in some P.1 cases) were observed. A two amino acid deletion at position 69-70 in the S protein is observed in cluster 5 and lineages B.1.1.7, B.1.525 and B.1.258 ^43^. The appearance of shared mutations in distinct and rapidly spreading SARS-CoV-2 lineages suggests that these mutations, either alone or in combination, may provide fitness advantage. In principle, mutations in the NTD, furin cleavage site or HR1 may affect the function of spike, as well as any mutation that results in conformational rearrangements of the RBD. Therefore, it is important to understand how synergetic interactions in spike collectively change the RBD.

Serum neutralizing antibodies were shown to develop upon natural SARS-CoV-2 infection ^44^ or vaccination ^45, 46^, and last at least several months ^47–49^. The alarm caused by the emergence of SARS-CoV-2 variants with the potential to escape host immunity acquired through infection or vaccination has been confirmed by a series of reports about lineages B.1.351 and P.1. ^31, 32, 36, 50, 51^. These raise concerns on whether a reduction in vaccine efficacy could result in re-infections and delay the reduction in mortality caused by circulating SARS-CoV-2. There is, therefore, the urgent need to develop mutation-tolerant vaccines (and biopharmaceuticals) targeting the skype protein. In this sense, it is critical to determine what makes mutations well tolerated for the viral lifecycle whilst efficiently escaping immunity. In this work, we engineered spike-pseudotyped viruses and analyzed individual and combined mutations that convergently appeared in different lineages, over time, and across several geographic locations, to determine their synergetic effects on neutralizing-antibody responses. We then used available structures of the complexes spike-antibodies and spike-ACE2 to determine the distance between each amino acid residue in the complex and the frequency of interactions of each amino acid residue of the RBD with either ACE2 or antibodies. We found a moderate reduction in neutralizing potency of sera against SARS-CoV-2 spike-pseudotyped lentivirus containing single mutations at position 484 (E484K) and 494 (S494P). Interestingly, the reduction became substantial with the addition of synergetic mutations K417N/N501Y or E484K/N501Y to E484K or S494P, respectively. In addition, we show that the amino acid residues at positions 484 and 494 frequently engage in binding to antibodies but not in binding to the receptor ACE2. Our work suggests that the amino acid residues at the RBD that are more dispensable for binding to ACE2 can more promptly evolve immune escape mutants if the amino acid residue substitution severely alters the binding specificity of the antibody.

## Results

### Construction of spike-pseudotyped viruses containing single or combined mutations in spike

Geographical distribution of mutations and associated prevalence is important, not only to track viral evolution and dynamics of lineages circulating worldwide, but also to identify recurrent mutations and ultimately tailor preventive measures to contain the virus. We used the pipeline https://github.com/wtsi-hpag/covidPileup [github.com] to trace single nucleotide polymorphisms (SNPs) in a given country or region. We explored a comprehensive data set containing 416,893 sequences of SARS-CoV-2 downloaded from GISAID to unravel the dispersal history and dynamics of spike mutations observed in SARS-CoV-2 viral lineages in the geographic locations in different continents divided as Australia (AUS), UK, European Union (EU), SA and USA, and integrated the data with viral circulation in these areas, measured by the number of new cases per week (fig. S1). Most regions were selected based on the appearance of specific lineages and variants of concern. Australia was included because of the restrictive measures of entering the country, closeness to Asia, geographical dispersion and for constituting an almost independent evolutionary landscape. These geographical locations were selected based on availability of sequenced samples during the period analyzed (from the 29^th^ of December 2019 until week 5 of 2021). Based on the reference genome NC_45512 (Wuhan-Hu-1, 29903 bases), the pipeline detected 3,766,497 SNPs and 2209 indels covering 25443 genome locations. For this study, we selected only mutations in spike because this protein is responsible for viral recognition of ACE2 receptor ^6, 12, 14, 52^ and for inducing neutralizing antibodies in the host ^10^. Of note, antibodies with neutralizing capacity were shown to bind to the NTD, but mostly to the RBD of spike ^2, 11, 50, 53, 54^. Many reports showed how single mutations change host neutralizing capacity and demonstrated that variants of concern B.1.351, P.1., or others including the mutation E484K, are able to escape immunity ^31, 35, 41, 55^. Whilst mutations on the RBD may sterically block binding of spike to ACE2, other mutations in spike may affect the conformation of the protein thereby impacting antibody recognition. Given this, we broaden our selection of mutations to include variants of concern and of interest (up to all mutations in lineages B.1.1.7, B.1.351, P.1., B.1.427/ B.1.429, table S1), high prevalence (fig. S2), or convergent appearance in different lineages over time. In terms of prevalence, the S477N mutation was accompanied by a peak of incidence in Australia from week 23-36 of 2020 and reached a prevalence of 20% worldwide (figs. S1 and S2). The L18F alone or combined with A222V was also accompanied by a peak in incidence in the UK and EU in weeks 38-45 of 2020, and in South Africa from week 42-to present and reached 20% incidence worldwide. Other single mutations include L452R and E484K, with a global prevalence of up to 6% worldwide; D839Y with peaks up to 2.5% during weeks 7-19 of 2020; Q675H, from week 23-present; deletion of amino acid residues at positions 69/70 combined with Y543H from week 39-51 of 2020; and D936Y from week 9-37 of 2020 (figs. S1 and S2). The phenotypic evaluation of SARS-CoV-2 mutations comprises several layers and includes the interaction of the virus with the host, disease severity and epidemiology. In this study, we evaluated evasion of host neutralizing antibodies.

All mutations engineered in this study, and the domain in spike in which they occur, are highlighted in Fig. 1B and C, detailed in table S1 and shown at the structural level in figs. S3-S5. In addition, to understand how interactions in spike collectively change the RBD, we evaluated individual or a combination of mutations on spike associated with variants of concern (up to all defining mutations, shown in table S1 and Fig. 2A-D) and have overall produced 25 different spike-pseudotyped viruses (listed in table S1). The mutations in the RBD region used in this study are highlighted also in the open (‘up’) conformation (Fig. 1C), showing the different conformational changes that each amino acid residue undergoes in this region.

**Fig. 1.**
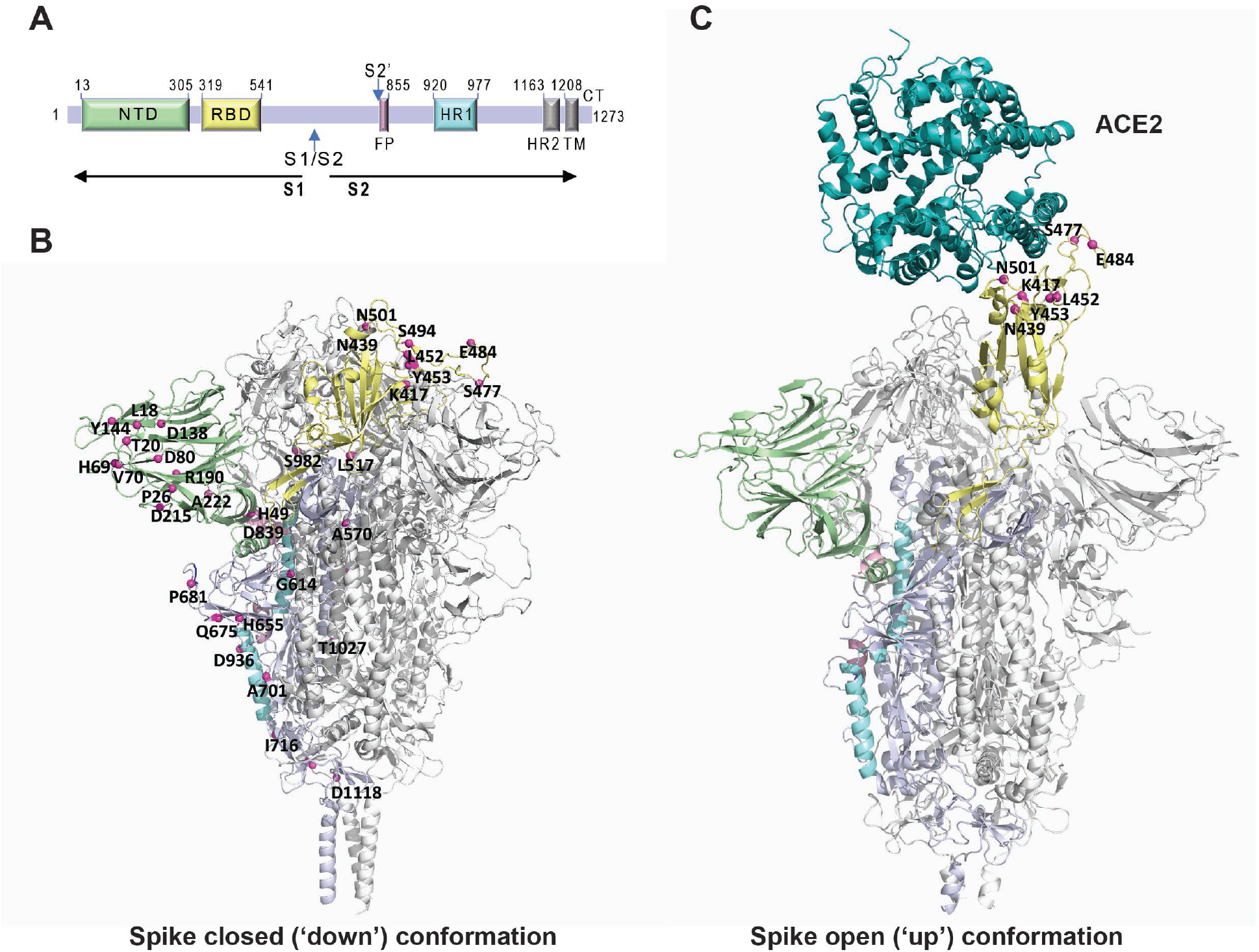
Location of the mutations tested in this work. **(A)** Schematic representation of the SARS-CoV-2 spike primary structure. NTD, N-terminal domain; RBD, receptor-binding domain; S1/S2, S1/S2 protease cleavage site; S2’, S2’ protease cleavage site; FP, fusion peptide; HR1, heptad repeat 1; HR2, heptad repeat 2; TM, transmembrane domain; CT, cytoplasmic tail. The protease cleavage sites are indicated by arrows. **(B)** The mutations in spike engineered in this study are mapped onto the spike protein surface, using a structure obtained by homology-based modelling with the structure PDB ID 6XR8 as a template. Mutations are highlighted using magenta spheres mapped on one of the monomers, which is represented with different colors to highlight important regions: NTD (green), RBD (yellow), FP (pink), HR1 (cyan). The remaining monomers are displayed in grey. **(C)** The structure of the spike protein with one RBD up and bound to ACE2 (PDB ID 7DF4) is shown to highlight the mutations used in this study located in the RBD in the context of its interaction with the receptor. The spike protein is represented in the same color scheme used in (A) and ACE2 is displayed in teal.

**Fig. 2.**
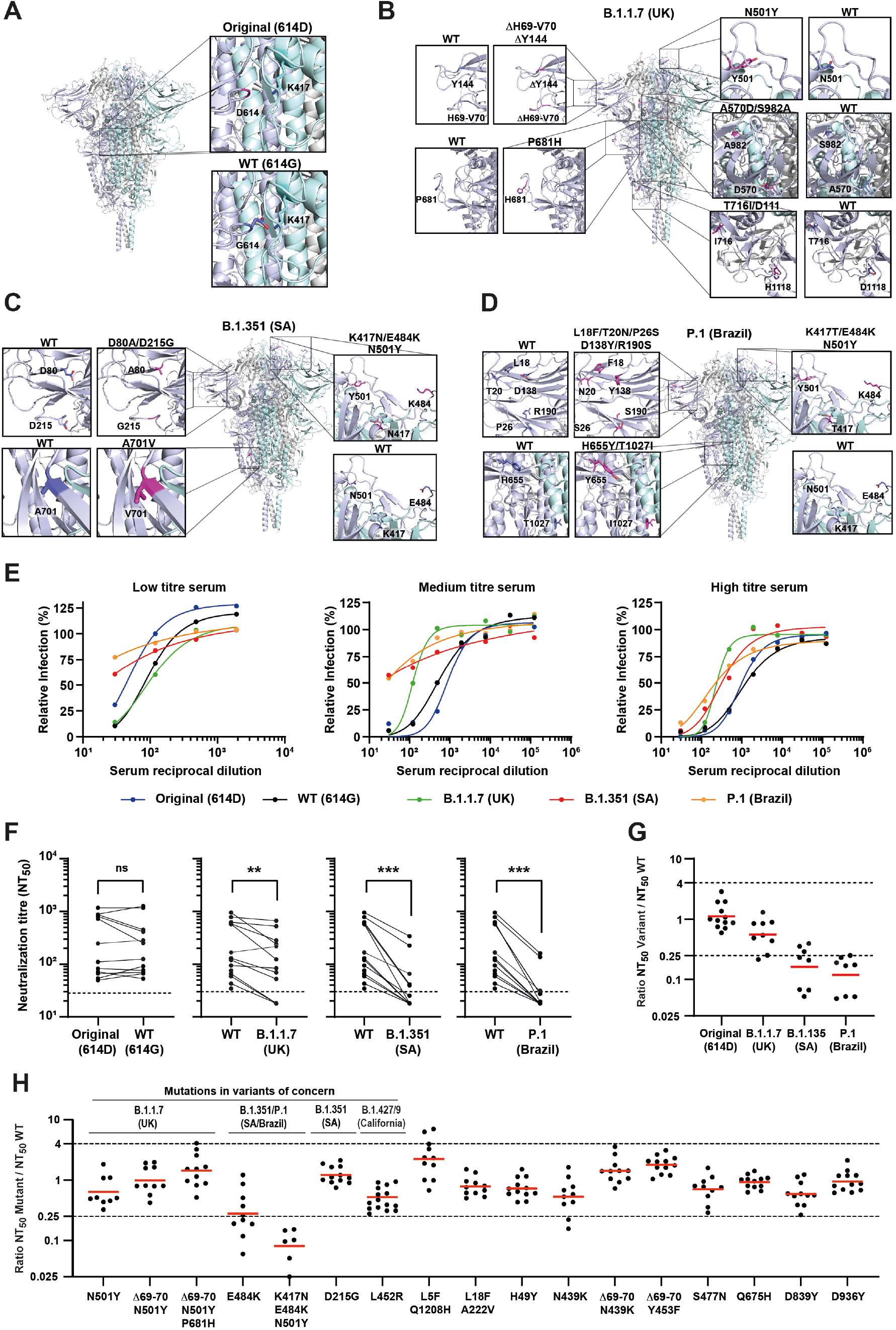
Neutralization of SARS-CoV-2 spike mutants by convalescent human sera. **(A-D)** SARS-CoV-2 spike secondary structure of the original strain (A), the B.1.1.7 variant first identified in the UK (B), the B.1.351 variant (SA) (C) and the P.1 variant (Brazil) (D). Insets show spike structure surrounding each mutated amino acid, and the corresponding structure in the WT. (**E-H)** Neutralization assays were performed by infecting 293T-ACE2 cells with pseudoviruses displaying WT or mutant SARS-CoV-2 spike, in the presence of serial dilutions of a convalescent human sera cohort (n=12-16). (**E**) Representative neutralization curves of sera with low, medium, and high titers of anti-spike IgG antibodies against WT virus, original virus, and variants of concern B.1.1.7, B.1.351 and P.1. Relative infection is defined as the percentage of infection relative to cells infected in the absence of serum. (**F)** Paired analysis of neutralizing activity of each serum against WT vs original virus or variants of concern. NT_50_ is defined as the inverse of the dilution that achieved a 50% reduction in infection. Dashed lines indicate the limit of detection of the assay (NT_50_=30). ns, non-significant, **p<0.01, ***p<0.001 by two-tailed Wilcoxon matched-pairs signed-rank test. (**G)** Ratios of neutralization between variant and WT viruses. (**H)** Ratios of neutralization between mutant viruses with single, double, or triple mutations and WT virus. Red bars indicate the geometric mean. See figs. S3-S5 for predictions of mutations in the structure of spike; S8 and S9 and table S2 for the complete set of neutralizing curves, neutralizing titers, anti-spike IgG titers determined by ELISA and the numerical values plotted in this figure.

The different spike versions served to engineer lentiviral spike-pseudotyped particles through a three-plasmid approach: a plasmid harboring the lentiviral genome, lacking Gag-Pol and envelope proteins, and encoding a GFP reporter; a Gag-Pol expression plasmid; and a plasmid expressing SARS-CoV-2 spike protein (or VSV-G protein, as control). Cells infected by these lentiviruses express GFP and can be easily quantified with high-throughput analytical methods.

### Neutralization assays on spike-pseudotyped viruses to evaluate synergetic effects of mutations

For the neutralization assay, we developed a 293T cell line expressing the human ACE2 receptor (fig. S6). The specificity of the assay was assessed using an anti-spike polyclonal antibody and an RBD peptide that competes with the spike protein for viral entry in SARS-CoV-2 spike-pseudotyped but not in VSV G-pseudotyped lentivirus that was used in parallel (fig. S7).

We characterized the serum antibodies from 12-16 health care workers, infected with SARS-CoV-2 during spring/summer 2020, for their ability to neutralize our spike-pseudotyped mutant lentiviruses. These sera were tested by ELISA for their anti-spike content and classified as low (up to endpoint titer 1:150), medium (endpoint titer 1:450) and high (endpoint titer from 1:1350) (table S2). We also used 4 negative sera as control (one pre-pandemic and 3 contemporary). Representative neutralization curves of the WT (614G) and original (614D) strains, as well as variants of concern B.1.1.7, B.1.351 and P.1 (containing all defining mutations, listed in table S1) are shown in Fig. 2E and the complete set of neutralization curves in fig. S8. For each neutralization curve, we calculated the half-maximal neutralization titer (NT_50_), defined as the reciprocal of the dilution at which infection was decreased by 50% (Fig. 2F and fig. S9). Consistent with the literature, the ELISA titer was associated with the neutralizing titer of sera (table S2) ^48, 56^.

In this paper, we used the Wilcoxon statistical test when comparing NT_50_s to understand if variants are significantly different from the wild-type. However, to define whether variants are capable of evading neutralization, we use the 4-fold criteria, similar to what has been done with influenza virus, where differences in hemagglutination assays greater than 4-fold highlight the need for vaccine update ^36, 57, 58^. Data show that, although the spike including all defining mutations of B.1.1.7 exhibited a modest reduction in neutralization titers relative to WT spike, it was still efficiently neutralized by convalescent sera, as observed before ^35, 59^. Conversely, the complete set of defining mutations of B.1.351 and P.1 consistently showed a reduction in neutralization capacity higher than 4-fold and are considered significantly different from the WT virus (Fig. 2G). We then analysed a series of single, double and triple mutations (Fig. 2H and figs. S6 and S7). We found that the mutations associated with B. 1.1.7 variant N501Y, Δ69-70/N501, Δ69-70/N501+P681H behave as WT spike. The mutation E484K alone reduced sera neutralization capacity by 3.6-fold [ratio NT_50_ mutant/NT_50_ WT of .028±0.37 (geometric mean ± SD)], and inclusion of additional mutations (K417N and N501Y) further decreased the neutralization (0.08±0.05), emphasizing the role of synergic mutations on immune evasion and underscoring the need for evaluating single and combined mutations on variants ^51^. Other mutations at the NTD, RBD, FP, HR1, TM and CD did not significantly alter sera neutralization efficiency. However, we observed a reduction in neutralization in some sera against N439K mutant (0.53±0.45) as reported in ^41^, which was not observed when Δ69-70 was added (Fig. 2H), and also against L452R mutant (0.48±0.23) that is associated with the most recent variants of concern B.1.427/9, as reported in ^60^. Importantly, we tested BNT162b2-elicited plasmas (collected after the first and second dosage of the Pfizer–BioNTech vaccine) for their capacity to neutralize the spike-pseudotyped particles expressing WT spike protein or mutants bearing all defining mutations of the variants of concern B.1.1.7, B.1.351 and P.1. Amongst 10 individuals, plasma collected 12 days after the administration of the first vaccine dose displayed anti-spike IgG titers from 1:450 to 1:12150 (table S3), and only one of them exhibited neutralizing capacity (table S3 and fig. S10). BNT162b2-elicited plasma collected 12 days after the administration of the 2^nd^ dose had very high levels of anti-spike IgG (titers 1:36450 −1:109350), higher than any sera collected upon natural infection, and all exhibited neutralizing activity against the variants of concern B. 1.351 and P.1, although with significantly lower efficiency than against the wild type or the B.1.1.7 variant.

Collectively, these data show that the spike-pseudotyped lentiviral particles used in this study, in which GFP is the output and is measured by high throughput and cheap methodologies, is suitable to quantitatively assess how single and synergetic mutations in full-length spike affect neutralizing-antibody responses.

### Frequency of contacts spike-antibodies and spike-ACE2

To determine which spike protein residues are more frequently targeted by neutralizing antibodies, we analyzed 57 structures retrieved from the protein data bank (PBD) containing the spike protein (or only the RBD region) bound to antibodies (table S4). Our results revealed that the receptor binding motif (RBM), in RBD, is the region with the highest frequency of contacts (Fig. 4A and B), consistent with what has been documented ^11, 61–64^. This region is important for the SARS-CoV-2 infection since it mediates contacts with ACE2 ^14, 65, 66^ and, for this reason, the binding of antibodies to this region would hinder ACE2 binding and, consequently, the entry of the virus in the cell.

In addition to contacts with the RBD, we also identified other regions of the spike protein with antibody binding sites. The antibodies FC05 and DH1050.1 bind to segments in the N-terminal region (between residues 143-152 and 246-257), region identified as an antigenic site ^54^, and 2G12 binds to residues 936-941 in the stalk (Fig. 4B and C).

To choose the cut-off above which to select the most important residues for antibody binding, we built a histogram (fig. S11). Three distinct groups appear: the first includes residues that have an interaction probability below 20%, the second includes residues whose interaction is between 20% and 45% and the last corresponds to frequent binders (interaction probability over 45%). Within this last group, we find Y449, L455, F456, E484, F486, N487, Y489, Q493, S494, and Y505 residues (Fig. 4A and B). More than half of these residues have hydrophobic side chains, showing that, within the batch of antibodies we studied, the most common binding mode is through hydrophobic contacts. These residues are important for antibody binding, which means that mutations within this group may enable the virus to escape antibodies from patients previously exposed to the WT or another variant, or people vaccinated using WT sequences of the S protein. In fact, prior studies have shown that mutations on most of these residues influence antibody escape ^2, 67^. For residues L455, F456 and F486, mutations for less hydrophobic residues have been shown to reduce binding by polyclonal antibodies ^67^ and mutants at site 487 escape human monoclonal antibodies COV2-2165 and COV2-2832(*2, 63*). Interestingly, our results also show that there is a high prevalence of antibodies binding to E484 (∼50%, Fig. 4B).

To be able to escape antibodies while maintaining the ability to efficiently infect cells, the mutations introduced must destabilize the interaction with antibodies while keeping a high binding affinity for ACE2. Thus, we reason that mutations occurring in residues that are relevant for antibody binding but not very relevant for the interaction with ACE2 can be advantageous for the virus. To gain further insights into this subject, we performed molecular dynamics (MD) simulations of the RBD bound to ACE2. These allowed us to determine which RBD residues are relevant for binding to ACE2, and the persistence of these interactions (Fig. 4D). Combining the information on antibody and ACE2 binding allows us to predict which are the most relevant amino acid residues (high affinity for antibodies and low affinity for ACE2). Using these criteria, two residues stand out: E484 and S494 (Fig. 4E, IV quadrant circled in red). These results are consistent with the evidence that E484 is an important mutation site and bring to light a new relevant site: S494. Mutations in S494 have been found in circulating strains, and the S494P mutation was shown to reduce the binding by polyclonal plasma antibodies ^67^ whilst having no ^4^ or modest effect ^68^ in RBD-ACE2 binding. To elucidate the role of this mutation in antibody neutralization, we used our neutralization assay and found that mutation S494P alone leads to a reduction in neutralization by covalence sera that is significant (Fig. 4F) despite not reaching our 4-fold criteria (ratio NT_50_ of mutant in relation to WT of 0.39±0.28, Fig. 4G). This value was further reduced when this mutation was combined with N501Y (0.28±0.21, Fig. 4G), or when combined with E484K (0.25±0.32, Fig. 4F-G). Interestingly, the E484K/S494P/N501Y triple mutant reduced neutralization over the 4-fold criteria applied (0.17±0.34, Fig. 4F-G). We also tested the ability of these mutants to evade neutralization by post-immunization sera, but the effect was less pronounced (Fig. 4H-I). In fact, S494P alone did not significantly reduce neutralization, becoming significant when N501Y (0.70±0.4), or E484K (0.5±0.3) mutations were introduced. The triple mutant displayed the more significant reduction in neutralization, despite not reaching the 4-fold difference (0.34±0.18, Fig. 4H-I). Importantly, S494P mutation appeared independently on several occasions and is under positive selection ^68^ and is now a variant in monitoring, especially when combined with E484K or N501Y ^23, 69^. Our results indicate that the S494P mutation can facilitate the escape of the virus from antibodies, maintaining the ability to bind to the receptor and enter host cells, and should be surveilled worldwide.

Some residues that are important for antibody binding are also involved in frequent interactions with ACE2. Seven of these residues remained bound to ACE2 throughout the whole simulation in all the replicates: L455, F456, F486, N487, Y489, Q493 and Y505 (Fig. 4D, top residues in quadrant II). This group includes residue F456, which was found to be a hotspot for escape mutations ^67^. Although this may seem contradictory, the same study shows that mutations in this residue were very rarely found in nature and, because of lack of prevalence, were not analyzed ^67^. We hypothesize that the low mutation rate for this site may be due to the high relevance of this residue for RBD-ACE2 binding. In fact, deep mutational analysis shows that when mutating F546 to any amino acid residue besides its SARS-CoV-1 counterpart (a leucine), the ACE2 binding affinity is reduced ^4^. Even though a viral strain with a single mutation at the site 456 could escape antibodies, it would be unfit to subsist in nature, due to a reduced ability to infect host cells.

Together, the analysis presented here is supported by experimental evidence on mutation-driven binding to ACE2 and escape to antibodies ^2–4^, as well was with the emergence of natural variants ^31, 35, 41, 55, 70–72^, and may be used to predict mutation hotspots.

## Discussion

It is critical to understand how SARS-CoV-2 will evolve and if it will escape host immunity, which could pose challenges to vaccination and render therapies ineffective.

We developed spike-pseudotyped lentiviral particles for high-throughput quantitation that express GFP upon entering cells, contributing to the toolkit of neutralizing assay methodologies ^73–75^. Our method is suitable to assess the neutralization activity of sera/plasma from individuals infected naturally or vaccinated, and to screen for antiviral drugs that block viral entry, such as therapeutic antibodies, in biosafety level 2 settings, which greatly facilitates the procedure and broadens its usage. It can be easily adapted to include single and multiple mutations, as observed in the present work for sera neutralizing activity (Figs. 2). Our results with BNT162b2-elicited plasma (Fig. 3) agree with previous publications showing resistance to neutralization by B.1.351 and P.1 lineages ^31, 32, 36, 50, 76^ and validate our neutralization assay. The fact that only mutations containing the E484K substitution promote immune escape, although this effect increases with synergic mutations that improve viral binding to ACE2 (K417N and N501Y), may be a consequence of lack in immunological selective pressure, a recognized driver of evolution ^58, 77–79^, as a large proportion of the population remains susceptible to SARS-CoV-2 infection. However, the emergence of this escape mutant suggests that the continued circulation of the virus may, in the future, impose further immunological constraints and result in viral evolution, as seen for influenza A virus ^58, 80^. How changes in SARS-CoV-2 will impact the circulation of the virus is not known, and the future will also elucidate whether vaccination will shape evolution of SARS-CoV-2 and permit reinfections. At the moment, reinfections are considered rare events ^81, 82^, but reports that SARS-CoV-2 escape mutants were shown to drive resurgence of cases upon natural infection ^26, 83^ are indicative that viral dynamics in the population may change ^29, 84^. In agreement, in this study and in reports by others, a reduction in neutralization activity by vaccination elicited plasma against variants of concern B.1.351 and P.1 was observed ^31, 32, 36, 50, 76^. Therefore, the development of methods able to predict hotspots of immune evasion are in demand. The surveillance of circulating strains across time, the evaluation of the type and duration of host immune responses upon natural infection and vaccination, and the identification of antibodies that efficiently control SARS-CoV-2 are essential measures to understand and control SARS-CoV-2 infection and host response ^85–87^. We argue that the structural information on complexes between antibodies, or ACE2, and spike variants may be integrated with mutational maps and their phenotypic characterization ^2–4^ to predict hotspots of immune evasion.

**Fig. 3.**
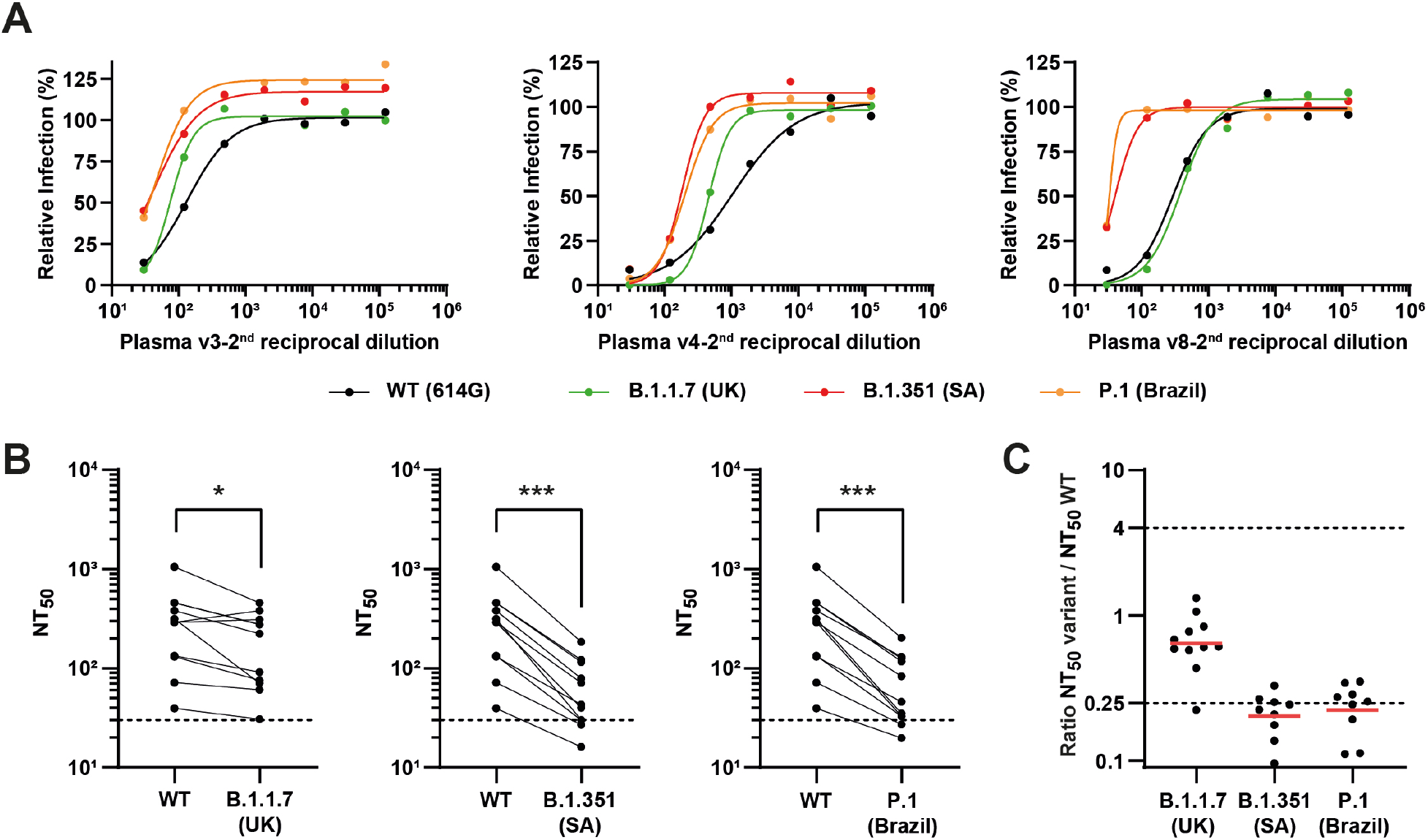
Neutralization of SARS-CoV-2 variants of concern by post-vaccination plasma. Neutralization assays were performed with pseudovirus displaying WT SARS-CoV-2 spike or variants of concern B.1.1.7, B.1.351 and P.1, in the presence of serial dilutions of post-vaccination plasmas from 10 individuals, 12 days after the first and the second rounds of vaccination. (**A)** Representative neutralization curves of post-vaccination plasma against WT virus and variants of concern**. (B)** Paired analysis of neutralizing activity each of plasma against WT vs variants of concern. Dashed lines indicate the limit of detection of the assay (NT_50_=30). *p<0.05, ***p<0.001 by two-tailed Wilcoxon matched-pairs signed-rank test. (**C)** Ratios of neutralization between variant and WT viruses. Red bars indicate the geometric mean. See fig. S10 table S3 for the complete set of neutralizing curves, the anti-spike IgG titers of each plasma and the numerical values plotted in this figure.

In this work, we analyzed by neutralization assays how a full-length spike carrying single or multiple mutations dispersed throughout the protein affected the conformation of the RBD. With this assay, we probe how antibodies from sera of infected patients or plasma collected from vaccinated people after administration of the 1^st^ and 2^nd^ dose of the vaccine block viral entry. We used structural information on the protein complexes spike-antibodies and spike-ACE2 to define the non-overlapping frequency of interactions using a cut-off value of 4.5 Å distance to define a contact. Given the idea that only a few amino acid residues are involved in protein-protein interfaces ^88^, including in antibody-antigen recognition, we aimed at identifying which amino acid residues in the RBD may evolve antibody escape mutants without affecting viral entry. With this approach, we identified two amino acid residues at the RBD-positions 484 and 494 (Fig. 4E) - that frequently engage in interactions with antibodies but not with ACE2. We observed that the E484 is relevant for antibody binding (Fig. 2G), in agreement with our and previous experimental results, and with the occurrence of this mutation in two variants that are becoming highly prevalent and were shown drive resurgence of infection in sites with high levels of seroconversion ^2, 36, 57, 71, 72, 83^. Additionally, we found that mutations in residue 494 may be problematic, either alone or combined with synergetic mutations, as it reduces neutralization competency of convalescent sera, and thus facilitates antibody escape without substantially altering the affinity for the ACE2 binding (Fig. 4E-G and fig. S12). The mutation S494P has been found in nature with a prevalence of 0.81% of all sequenced genomes (week 11 of 2021, fig. S2) and has already been observed in combination with mutation E484K (reported on the 22^nd^ October 2020). Of note, current prevalence of B.1.351 and of P.1 is 1.9% and 0.47%, respectively, being in the same range as S494P. We posit that the prevalence of mutations in amino acids 484, 494 and 501 will increase with viral circulation and/or spike seroconversion and should be surveilled worldwide. We observed a similar pattern relative to the substitution L454R/Q in India, in which the E484K mutation was acquired posteriorly. At the moment, however, with the majority of the population still susceptible to viral infection, mutations that increase viral transmission, such as N501Y ^84^, have a selective advantage.

**Fig. 4.**
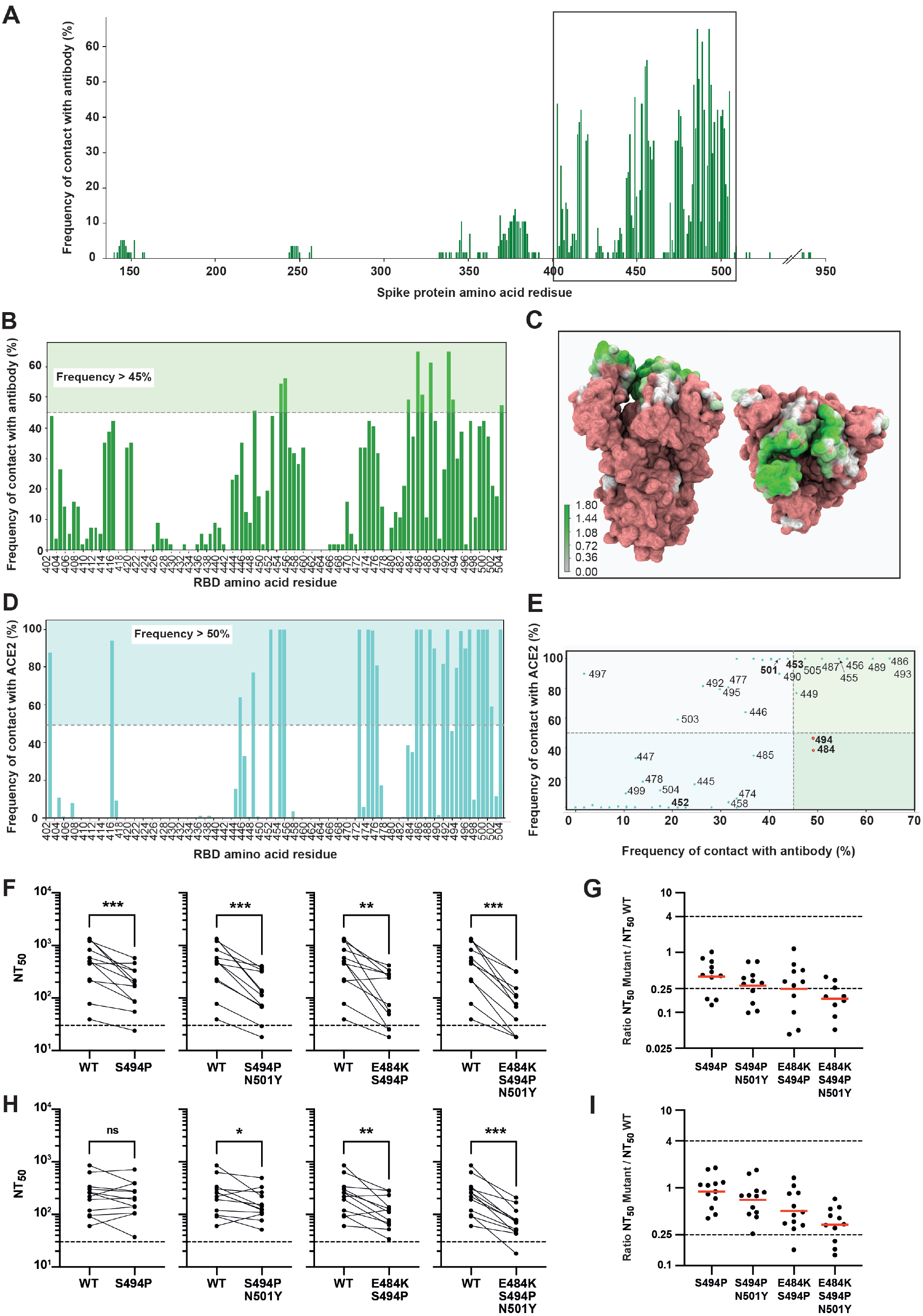
Spike protein-antibody and RBD-ACE2 contact analysis. **(A)** Histogram of the contact frequency of each spike residue with antibodies using a cut-off value 4.5 Å to define a contact. **(B)** Zoom in of the histogram shown in (A) in the region comprising the residues with the highest frequency of antibody contacts (located in the RBD). **(C)** Antibody contact map represented on the Spike protein surface, seen from front (left side) and from top (right side); the map was built with VMD ^108^ and the color gradient is used to represent the log of the frequency of contact with Ab as shown in the scale bar. **(D)** Histogram of the contact frequency of RBD residues with ACE2, obtained from MD simulations, using a cut-off value 4.5 Å to define a contact. **(E)** Scatter plot correlating the RBD residues’ frequency of contact with ACE2 and antibodies; each point represents one residue and those that were studied experimentally are highlighted in bold; the quadrants are filled with different colors to highlight the relevance of different residues as potential mutation hotspots, with the most relevant residues located in quadrant IV and shown as red points. **(F-I)** Neutralization assays were performed against pseudovirus displaying WT or S494P spike mutants, in the presence of serial dilutions of convalescent sera (F, G) or post-vaccination sera 1 and 30 days after the second round of vaccination (H, I). **(F, H)** Paired analysis of neutralizing activity of convalescent sera (F) or post-vaccination sera (H) against WT vs S494P, S494P/N501Y, E484K/S494P or E484K/S494P/N501Y mutants. Dashed lines indicate the limit of detection of the assay (NT_50_=30). ns, non-significant, *p<0.05, **p<0.01, ***p<0.001 by two-tailed Wilcoxon matched-pairs signed-rank test. (**G, I)** Ratios of sera neutralization between S494P mutants and WT viruses. Red bars indicate the geometric mean. See figs. S11-S12 for protein-antibody contact frequency histogram, overall representation of amino acid residues relevant for antibody and ACE2 binding and list of references of the structures of the antibodies used, respectively; figs. S13-S14 and tables S5-S6 for the complete set of neutralizing curves, the anti-spike IgG titers of each serum and the numerical values plotted in this figure.

Our approach has some caveats. Regarding the neutralization assays, and despite spike being the immunodominant protein targeted by antibodies ^7–11^, other viral proteins may contribute, even if in a small proportion to neutralization activity *in vivo*, which we are unable to detect using spike-pseudotyped particles. In addition, the sera/plasma we used was obtained at a fixed time interval. Future experiments should repeat the analysis using sera/plasma of people infected with known genotypes, or from different geographical areas and obtained at different times along the epidemics dynamic. Finally, we inspected a handful of mutations and not a comprehensive mutational map across spike ^2, 4^. However, we traced several mutations across spike in single and multiple combinations, and thus, we conclude that our analysis is relevant to inform on mutation-driven escape viruses to antibodies. The analysis of frequency of interactions between spike-antibodies or ACE2 has some issues associated as well. The pool of complexes spike protein/RBD and antibodies analyzed here is not necessarily representative of the universe of antibodies found in COVID-19 patients, since it is limited by the availability of structural information and is biased to monoclonal antibodies (table S4). All the antibodies analyzed are neutralizing antibodies, which is not the case in nature ^54^. Additionally, many structural studies have focused on the interaction with the RBD, while other regions have not been so exhaustively analyzed, which creates a bias in terms of the number of RBD binding antibodies with known structure ^14, 65, 66^. Limitations of the methods used, such as crystallographic contacts, can also influence the results. Nevertheless, its combination with data from the molecular simulation and from experiments helps to predict and explain the emergence of new mutations.

In conclusion, our analysis of the spike protein-antibody contacts revealed that (within the available data set) the receptor binding motif (RBM), in the receptor binding domain (RBD), is an important region for antibody binding, which makes sense, since antibody binding to this region can prevent the interaction with ACE2 and ultimately impair viral infection. Our results are in very good agreement with experimental data on mutation-driven escape to polyclonal antibodies and with the emergence of natural variants, which were reported to re-circulate in previously exposed people ^83^. Thus, this analysis represents a simple strategy to predict mutation hotspots that should be kept under surveillance. The predictions will become even better once the pool and diversity of available structures increases.

## Methods

### Usage of COVID-19 pipeline for geographical SARS-CoV-2 variation analysis

We have used a pipeline for COVID-19 variation analysis using whole genome sequences. CovidPileup can be downloaded from https://github.com/wtsi-hpag/covidPileup. The pipeline includes SNP and indel calling and tracing specific SNPs in a given country or region. By incorporating metadata, it is also possible to assign tags such as collecting time, age and sex to each identified SNP or indel. To allow quick search and data processing, a multi-layered indexed data structure has been designed. We assume the size of the reference is G and the number of sequenced samples is N. The first index implemented is the genome reference coordinates which points to SNP or indel locations while the second index stores the number of SNPs and the third index records the number of countries SNPs are associated with. When a search is pursued on a location, one can quickly get the information on the numbers of samples containing variation and countries on this location. Specifically, if all the SNPs are from one country or one region (such as EUR or UK), the variant will be a specific or unique SNP/indel. Statistics of COVID-19 variations was performed on CovidPileup files, based on weeks (week 1, Dec 29th 2019 - Jan 5th 2020). Some samples with reported problems were excluded, according to the publicly available document from nextstrain https://github.com/nextstrain/ncov/blob/master/defaults/exclude.txt. Data for new positive cases per week is based on WHO Coronavirus Dashboard (https://covid19.who.int/WHO-COVID-19-global-data.csv). The following regions of world were defined: EUR (EU region, samples with tag ‘France’, ‘Germany’, ‘Italy’, ‘Belgium’, ‘Netherlands’, ‘Austria’, ‘Denmark’, ‘Sweden’, ‘Norway’, ‘Poland’, ‘Portugal’, ‘Switzerland’, ‘CzechRepublic’, ‘Iceland’, ‘Luxembourg’, ‘Greece’, ‘Hungary’, ‘Slovenia’, ‘Slovakia’, ‘Malta’, ‘Croatia’, ‘Slovenia’, ‘Latvia’, ‘Lithuania’, ‘Estonia’, ‘Bulgaria’, ‘Cyprus’ and ‘Ireland’), UK (Samples with tag ‘UnitedKingdom’, ‘England’,’NorthernIreland’, ‘Scotland’, and ‘Wales’), Australia, USA and South Africa. Prevalence of amino acid residue genotypes of spike proteins were counted and shown in fig. S1. Minor genotypes with < 1% prevalence or supported by < 3 samples were grouped as “other genotype”.

### Cells and plasmids

Human hepatocellular carcinoma HuH-7 cells (a kind gift from Dr Colin Adrain, Instituto Gulbenkian de Ciência, Portugal), Human Embryonic Kidney 293T (provided by Prof Paul Digard, Roslin Institute, UK) and 293ET cells (from Dr Colin Adrain) were maintained in Dulbecco’s Modified Eagle Medium (DMEM, Gibco, 21969035) supplemented with 10 % fetal bovine serum (FBS, Gibco, 10500064), 1% penicillin/streptomycin solution (Biowest, L0022) and 2mM L-glutamine (ThermoFisher, 25030024), at 37°C and 5% CO2 atmosphere. Plasmids used in this study were obtained as follows: pLEX.MSC was purchased from Thermo Scientific; psPAX2 and pVSV.G were a kind gift from Dr Luís Moita (Instituto Gulbenkian de Ciência, Portugal); pEGFP-N1 was provided by Dr Colin Crump (University of Cambridge, UK); and pCAGGS containing the SARS-Related Coronavirus 2, Wuhan-Hu-1 Spike Glycoprotein Gene (pCAGGS-SARS-CoV-2-S, NR-5231) was obtained through BEI Resources.

### Cloning and S protein mutations

To facilitate incorporation of S protein into lentiviral pseudovirions, the C-terminal 19 amino acids encompassing the ER-retention domain of the cytoplasmic tail of S protein were deleted. This construct, named SARS-CoV-2 S_trunc_, was generated by PCR amplification of the C-terminal region of S without amino acids 1255-1273 (a premature STOP codon was added in the reverse primer), using pCAGGS-SARS-CoV-2-S as template (primers in table S7). The amplified sequence was cloned back into pCAGGS-SARS-CoV-2-S, between *Age*I and *Sac*I restriction sites, generating the expression vector pCAGGS-SARS-CoV-2-S_trunc_. Mutations were introduced in the spike protein by site-directed mutagenesis, with the Quickchange Multi Site-Directed Mutagenesis kit (Agilent) following the manufacturer’s instructions (primers in table S7). Lentiviral reporter plasmid pLEX-GFP was produced by PCR amplifying GFP from pEGFP-N1 (primers in Table 1) and cloning the insert into *BamH*I–*Xho*I restriction sites in the multi-cloning site of pLEX.MCS vector. For lentiviral plasmid pLEX-ACE2 production, human ACE2 coding sequence was amplified from Huh7 cDNA and cloned into pLEX.MCS, using *Xho*I and *Mlu*I restriction sites (primers in table S7).

### Production of 293T cells stably expressing human ACE2 receptor

To produce VSV-G pseudotyped lentiviruses encoding the human ACE2, 293ET cells were transfected with pVSV-G, psPAX2 and pLEX-ACE2 using jetPRIME (Polyplus), according to manufacturer’s instructions. Lentiviral particles in the supernatant were collected after 3 days and were used to transduce 293T cells. Three days after transduction, puromycin (Merck, 540411) was added to the medium, to a final concentration of 2.5 μg/ml, to select for infected cells. Puromycin selection was maintained until all cells in the control plate died and then reduced to half. The 293T-Ace2 cell line was passaged six times before use and kept in culture medium supplemented with 1.25 μg/ml puromycin. ACE2 expression was evaluated by flow cytometry (fig. S6).

### Production and titration of spike pseudotyped lentiviral particles

To generate spike pseudotyped lentiviral particles, 3×10^6^ 293ET cells were co-transfected with 8.89ug pLex-GFP reporter, 6.67ug psPAX2, and 4.44ug pCAGGS-SARS-CoV-2-S WT or mutants (or pVSV.G, as a control), using jetPRIME according to manufacturer’s instructions. The virus-containing supernatant was collected after 3 days, concentrated 10 to 20-fold using Lenti-XTM Concentrator (Takara, 631231), aliquoted and stored at −80°C. Pseudovirus stocks were titrated by serial dilution and transduction of 293T-Ace2 cells. At 24h post transduction, the percentage of GFP positive cells was determined by flow cytometry, and the number of transduction units per mL was calculated.

### Flow cytometry

Flow cytometry was performed as in ^89^. In brief, HEK 293T and 293T-ACE2 cells were prepared for flow cytometry analysis by detaching from the wells with trypsin, followed by fixation with 4% paraformaldehyde). For analysis of ACE2 expression, cells were stained with a primary antibody against ACE2 (4µg/ml, R&D Systems, catalog no. AF933) followed by a secondary antibody labelled with Alexa Fluor 568 (1:1000; Life Technologies). Analysis of cell populations was performed in a Becton Dickinson (BD) LSR Fortessa X-20 equipped with FACS Diva and FlowJo (BD, Franklin Lakes, NJ) software’s.

### Human convalescent sera and post-vaccination plasma/serum

Venous blood was collected by standard phlebotomy from health care providers who contracted COVID-19 during spring/summer 2020, as tested by RT-PCR on nasopharyngeal swabs. All participants provided informed consent to take part in the study. This study “Fatores de susceptibilidade genética e proteçao imunológica à COVID-19” was approved on the 25^th^ of May 2020 by the Ethics committee of the Centro Hospitalar Lisboa Ocidental, in compliance with the Declaration of Helsinki, and follows international and national guidelines for health data protection. Serum was prepared using standard methodology and stored at −20°C.

Peripheral blood from health care workers was collected by venipuncture into EDTA tubes at day 12 and day 33 post-immunization with Pfizer BTN162b2 mRNA vaccine and immediately processed. Alternatively, post-vaccination serum was collected at day 22 and day 51 post-immunization with Pfizer BTN162b2 mRNA vaccine. All participants provided informed consent and all procedures were approved by the ethics committee of Centro Hospitalar Lisboa Central, in accordance with the provisions of the Declaration of Helsinki and the Good Clinical Practice guidelines of the International Conference on Harmonization. Peripheral blood was diluted in PBS 1x (VWR), layered on top of biocoll (Biowest) and centrifuged at 1200g for 30 min without break. Plasma was collected to cryotubes and stored at −80°C ultra-low freezer until subsequent analysis.

### ELISA assay

Direct ELISA was used to quantify IgG anti-full-length Spike in convalescent sera. The antigen was produced as described in ^90^. The assay was adapted from ^91^ and semi-automized to measure IgG in a 384-well format, according to a protocol to be detailed elsewhere. For titer estimation, sera were serially diluted 3-fold starting in a 1:50 dilution and cut off was defined by pre-pandemic sera (mean + 2 standard deviation).

ELISA assay on post-vaccination plasma and serum was performed based on the protocol ^92^ and modified as described in Gonçalves *et al.* ^93^. Briefly, 96 well plates (Nunc) were coated with 50 µl of trimeric spike protein at 0.5 µg/mL and incubated overnight at 4°C. On the following day, plates were washed three times with 0.1% PBS/Tween20 (PBST) using an automatic plate washer (ThermoScientific). Plates were blocked with 3% bovine serum albumin (BSA) diluted in 0.05% PBS/T and incubated 1h at room temperature. Samples were diluted using 3-fold dilution series starting at 1:50 and ending at 1:10,9350 in 1% BSA-PBST/T and incubated 1h at room temperature. Plates were washed three times as previously and goat anti-human IgG-HRP secondary antibodies (Abcam, ab97215) were added at 1:25,000 and incubated 30 min at room temperature. Plates were washed three times and incubated ∼7min with 50 µl of TMB substrate (BioLegend). The reaction was stopped with 25µl of 1M phosphoric acid (Sigma) and read at 450nm on a plate reader (BioTek). Each plate contained 6 calibrator samples from two high-, two medium-, and two low-antibody producer from adult individuals collected at Hospital Fernando Fonseca that were confirmed positive for SARS-CoV-2 by RT-PCR from nasopharyngeal and/or oropharyngeal swabs in a laboratory certified by the Portuguese National Health Authorities ^93^. As negative control, we used pre-pandemic plasma samples obtained from healthy donors collected prior July 2019. The endpoint titer was defined as the last dilution before the absorbance dropped below OD_450_ of 0.15.

### Neutralization assay

Heat-inactivated serum and plasma samples were four-fold serially diluted over 7 dilutions, beginning with a 1:30 initial dilution. Dilutions were then incubated with spike pseudotyped lentiviral particles for 1h at 37°C. The mix was added to a pre-seeded 96 well plate of 293T-ACE2 cells, with a final MOI of 0.2. At 48h post-transduction, the fluorescent signal was measured using the GloMax Explorer System (Promega). The relative fluorescence units were normalized to those derived from the virus control wells (cells infected in the absence of plasma or serum), after subtraction of the background in the control groups with cells only. The half-maximal neutralization titer (NT_50_), defined as the reciprocal of the dilution at which infection was decreased by 50%, was determined using four-parameter nonlinear regression (least squares regression without weighting; constraints: bottom=0) (GraphPad Prism 9). To assess the specificity of our assay, an anti-Spike antibody neutralization assay and an RBD competition assay were performed in SARS-CoV-2 S pseudotyped and vesicular stomatitis virus (VSV) G pseudotyped lentiviral particles, in parallel (fig. S9). The anti-Spike antibody neutralization assay was performed as above, starting with a concentration of 50µg/ml (Thermo Fisher Scientific, PA5-81795) and three-fold serially diluting over 7 dilutions. For the RBD competition assay, 293T-ACE2 cells were pre-incubated with three-fold serial dilutions of SARS-CoV-2 spike’s receptor binding domain, starting with a concentration of 400 µg/ml. After 1h incubation at 37°C, supernatant was aspirated and pseudoviruses were added at an MOI of 0.2. The half maximal inhibitory concentration (IC_50_) was determined using four-parameter nonlinear regression (least squares regression without weighting; constraints: bottom=0) (GraphPad Prism 9).

### Spike protein-antibody contact analysis

In order to determine which residues of the SARS-CoV-2 spike protein contribute the most for the binding of antibodies, we studied 57 PDB structures containing the S protein (or only the RBD region) bound to antibodies (Table S1). These complexes were chosen based on their availability in the PDB repositorium ^94^ and are all neutralizing antibodies. We first used the MDAnalysis library ^95, 96^ to identify the residues of the antibody and the spike protein located in the interface between the two proteins. This was done for all PDB structures using in-house Python scripts. To determine the relevance of each spike protein/RBD residue in the binding, a distance cut-off value of 4.5 Å was applied as a criterion (i.e., a contact is observed when two residues have a minimum distance lower than 4.5 Å). Finally, the numerical python library (NumPy ^97^) was used to calculate the frequency of contact for each spike protein residue with an antibody residue, from the sample of PDB structures analyzed.

### MD simulation of the RBD-ACE2 complex

Molecular dynamics (MD) simulations of the RBD bound to the ACE2 protein were performed with the GROMACS 2020.3 package ^98^, using the Amber14sb ^99^ force field, starting from the 6m0j structure ^12^, in a truncated dodecahedron box filled with water molecules (minimum of 1.2 nm between protein and box walls). The TIP3P water model ^100^ was used and the total charge of the system (−23, including the constitutive Zn^-^ and Cl^-^ ions bound to ACE2) was neutralized with 23 Na^+^ ions. Additional Na^+^ and Cl^+^ ions were added to the solution to reach an ionic strength of 0.1M. The system was energy-minimized using the steepest descent method for a maximum of 50000 steps using position restraints on the heteroatom positions by restraining them to the crystallographic coordinates using a force constant of 1000 kJ/mol in the X, Y and Z positions. Before performing the production runs, an initialization process was carried out in 5 stages of 100 ps each. Initially, all heavy-atoms were restrained using a force constant of 1000 kJ/mol/nm, and at the final stage only the only C-α atoms were position-restrained using the same force constant. In the first stage, the Berendsen temperature algorithm ^101^ was used to initialise and maintain the simulation at 300 K, using a temperature coupling constant of 0.01 ps, without pressure control. The second stage continues to use the Berendsen temperature algorithm but with a force constant of 0.1 ps. The third stage kept the same temperature control settings, but introduced pressure coupling with the Berendsen pressure algorithm ^101^ with a pressure coupling constant of 5.0 ps and applied isotropically. The fourth stage changed the temperature algorithm to V-rescale ^102^, with a temperature coupling constant of 0.1 ps, and the pressure algorithm to Parrinello-Rahman ^103^ with a pressure coupling constant of 5.0 ps. The fifth stage is equal to the fourth stage, but position restraints are only applied to C-α atoms. During the simulation, the equations of motion were integrated using a timestep of 2 fs. The temperature was maintained at 300 K, using the V-rescale ^102^ algorithm with a time constant of 0.1 ps, and the pressure was maintained at 1 bar using the Parrinello−Rahman ^103^ pressure coupling algorithm, with a time constant of 5 ps; pressure coupling was applied isotropically. Long-range electrostatic interactions were treated with the PME ^104, 105^ method, using a grid spacing of 0.12 nm, with a cubic interpolation. The neighbor list was updated every twenty steps and the cutoff scheme used was Verlet with 0.8nm as the real space cut-off radius. All bonds were constrained using the LINCS algorithm ^106^. The system was simulated for 8 µs in 5 replicates (to a total of 40 µs).

### RBD-ACE2 contacts frequency

In order to determine the residues that contribute the most for the interaction between RBD and ACE2, and the persistence of these interactions, we performed a contact analysis throughout the simulation. We started by eliminating the first equilibration µs of all replicates. The MDAnalysis library ^95, 96^ was then used to pinpoint the residues of the RBD that are in contact with the ACE2 protein. A distance cut-off value of 4.5 Å was applied as a criterion (i.e., a contact is observed when two residues have a minimum distance lower than 4.5 Å). Finally, we determined the percentage of time for which a given RBD residue is at less than 4.5 Å of ACE2.

### Homology-based models of S protein variants

Homology-based models of all the variants analyzed in this study were generated using the software Modeller ^107^, version 9.23, using the structure of the wild type enzyme (PDB code: 6XR8) ^15^ as a template. The protocol used only optimizes the atoms belonging to the mutated residues and the residues that are located within a 5 Å radius from these residues, maintaining the remaining atoms fixed with the coordinates found in the template structure. The optimization parameters were set to the software default values. The final model corresponds to the one with the lowest value of the DOPE function out of 20 generated structures.

## Acknowledgments

The authors acknowledge Sean Whelan (Washington University School of Medicine St Louis) for providing reagents, João F. Viana and his team for organizing sera collection at Centro Hospitalar Lisboa Ocidental in Portugal, and the IGC COVID-19 team (Patrícia C. Borges, Christian Diwo, Paula Matoso, Vanessa Malheiro, Lígia A. Gonçalves, Nádia Duarte, Ana Brennand, Lindsay Kosack and Onome Akpogheneta) for processing and ranking samples, for this paper and during early settings. We are grateful to the Unit of Genomics of the Instituto Gulbenkian de Ciência for technical support, sample processing and data collection. The authors thank Isabel Gordo and Mónica Dias (Instituto Gulbenkian de Ciência) for helpful discussions andRute Castro and Paula M. Alves (Instituto de Biologia Experimental e Tecnológica, Universidade Nova de Lisboa, Portugal) for providing the antigen used in the ELISA and in the RBD competition assay.

## Author contributions

MJA and CS conceived and supervised the study. All authors designed and conducted experiments; all authors carried out experiments and analyzed the data; MJA, MA, RL, CMS, DL, MV wrote the manuscript; all authors contributed to editing the manuscript.

## Data and materials availability

All data are available in the main text or the supplementary materials.

## Supplementary Materials

**Figure S1:**
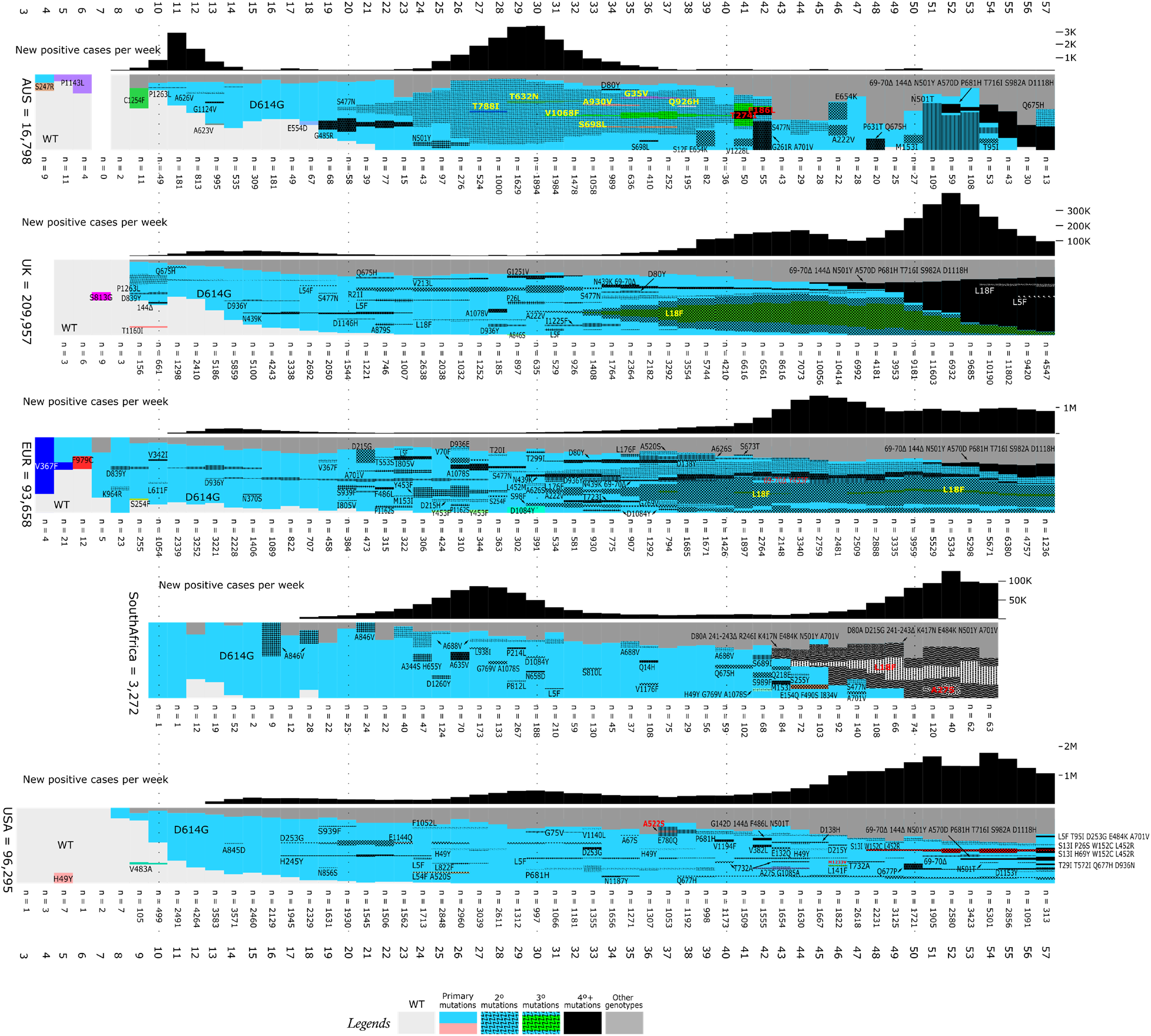
Geographical spike mutations and prevalence of COVID-19. Upper part each figure, new COVID-19 positive cases per week. Lower part, prevalence of spike mutations. Regions are highlighted on the left side as well as number of sequences analysed. The following regions of world were defined: Australia, UK (GISAID Samples with tag ‘United Kingdom’, ‘England’, ‘Northern Ireland’, ‘Scotland’, and ‘Wales’), EUR (Samples with tag ‘France’, ‘Germany’, ‘Italy’, ‘Belgium’, ‘Netherlands’, ‘Austria’, ‘Denmark’, ‘Sweden’, ‘Norway’, ‘Poland’, ‘Portugal’, ‘Switzerland’, ‘Czech Republic’, ‘Iceland’, ‘Luxembourg’, ‘Greece’, ‘Hungary’, ‘Slovenia’, ‘Slovakia’, ‘Malta’, ‘Croatia’, ‘Slovenia’, ‘Latvia’, ‘Lithuania’, ‘Estonia’, ‘Bulgaria’, ‘Cyprus’ and ‘Ireland’), South Africa and USA. Light grey - wild type (NC_045512 reference). Other genotypes are shown on the figure by shadows or colours. Dark grey regions are minor genotypes either < 1% prevalence or supported by < 3 samples.

**Fig. S2.**
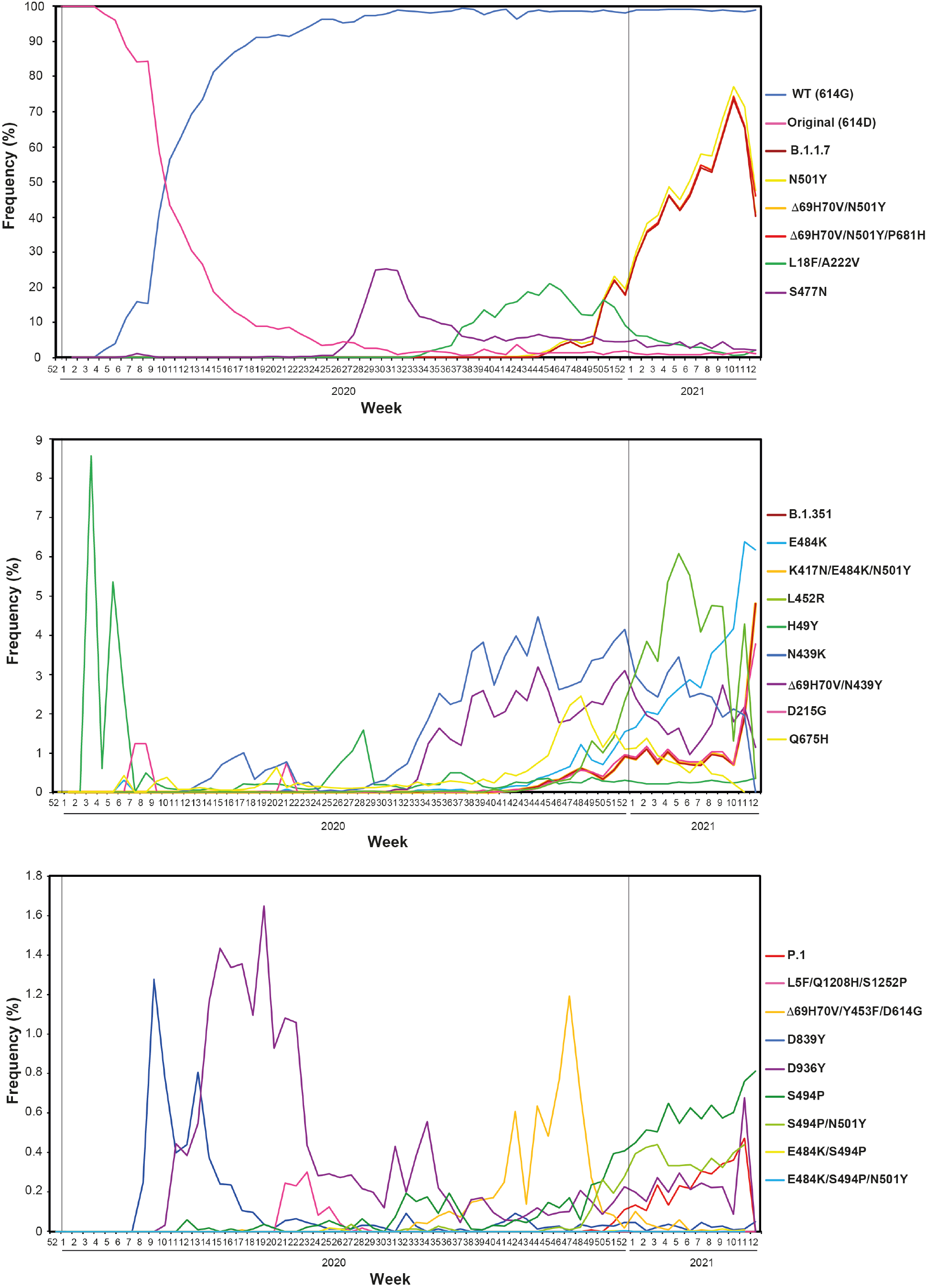
Temporal worldwide prevalence of spike mutations considered in this study from the 1^st^ January 2020 to the end of March 2021. In order to investigate the global frequency of SARS-CoV-2, the variant surveillance dataset was retrieved on March 25th, 2021 from GISAID (Shu Y, McCauley J., 2017). From this dataset and using text mining tools, the number and frequency of predominant variants was determined from Spike protein mutations and displayed per weeks according to variant collection date. The mutations are divided onto 3 graphs in which frequencies are up to 100% (upper graph), 9% (middle graph) and 2.5% (lower graph) for easier visualization. The frequencies are not cumulative but individual representations of each mutation. Note that week 11 and 12 of 2021 have had a reduction in the number of sequenced genomes, which is reflected in an overall reduction of circulating variants.

**Fig. S3.**
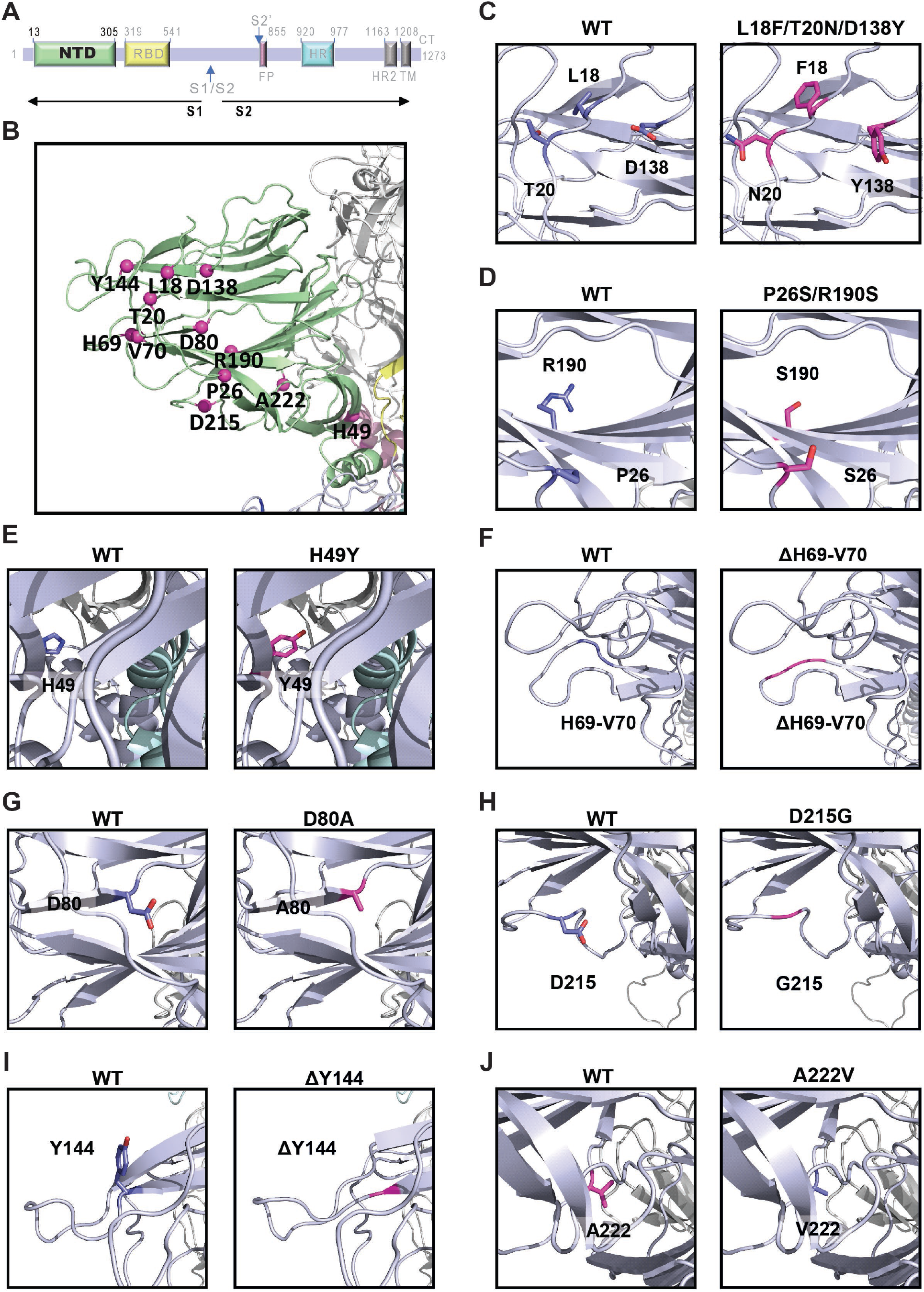
Mutations in the N-terminal domain of spike. **(A)** Schematic representation of the SARS-CoV-2 spike primary structure. **(B)** The mutations in spike NTD engineered in this study are mapped onto the spike protein surface. Mutations are highlighted using magenta spheres. **(C)** Residues 18 and 20 are in an exterior loop. The mutation to L18F enables a pi-stacking interaction with F127, which in the WT also interacts with L18 through hydrophobic contacts. The mutation T20N is not predicted to have a strong impact since T20 does not form specific interactions. Residue D138 sits in a semi-exposed loop and forms a salt bridge with R21, which is abolished when this residue is mutated to Y. **(D)** Residue 26 is in an exterior loop. Changing P26 to S might make the end of this loop more flexible, since prolines tend to confer rigidity to proteins. R190 is in a beta-sheet and can interact with H207 through pi-stacking, thus the mutation to S may destabilize this interaction and the secondary structure. **(E)** H49 is in an interior beta-sheet and interacts with R44 through pi-stacking, which is maintained upon the mutation to Y. **(F)** H69 and V70 are in a loop that has the appropriate features to be targeted by antibodies, since it is exposed and moderately hydrophobic. If this is the case, the deletion of these residues may impact the interaction with antibodies. **(G)** D80 is located at the tip of a semi-exposed beta-strand and forms a salt bridge with R78, which is lost with the mutation to A. **(H)** D215 sits in an exposed loop and forms a hydrogen bond with T29 that disappears with the D215G mutation, which may also make this loop more flexible. **(I)** Y144 sits at the tip of an exposed beta-strand and its deletion may impact the secondary structure and the possible recognition by antibodies **(J)** A222 is in a hydrophobic cleft and the mutation to V strengthens hydrophobic contacts. One additional mutation, L5F, has been engineered in combination with Q1208H. However, this mutation located to unstructured regions and hence its structure is unknown.

**Fig. S4.**
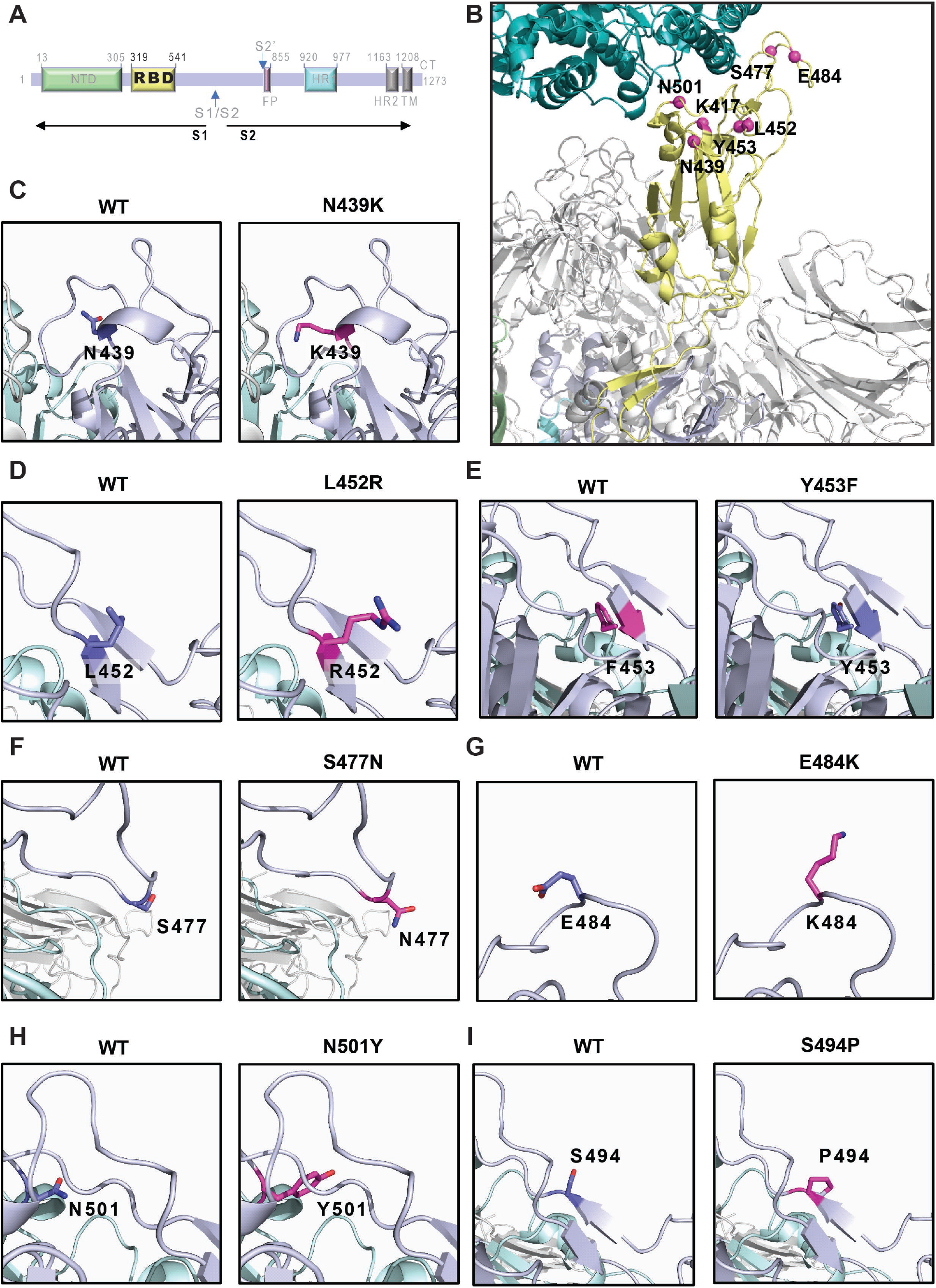
Mutations in the receptor binding domain of spike. **(A)** Schematic representation of the SARS-CoV-2 spike primary structure. **(B)** The mutations in spike RBD engineered in this study are mapped onto the spike protein surface. Mutations are highlighted using magenta spheres. **(C)** N439 sits in an exposed helix turn of the RBD and forms a hydrogen bond with S45, which is predicted to be lost with the mutation to K. This residue is not involved in RBD binding to ACE2. **(D)** L452 is at the tip of a short-exposed beta-strand of the RBD and does not interact with ACE2. Its mutation from hydrophobic L to the positively charged residue R may lead to changes in antibody interaction, since it is exposed. **(E)** Y453 is in the same B-sheet has L453 and interacts with H34 from ACE2 through pi-stacking, which can be maintained in the mutant Y453R. **(F)** S477 is in an exposed loop of the RBD, which often interacts with antibodies. S477 does not interact with ACE2. The mutation S477N is not predicted to have a strong effect since both residues are polar. **(G)** E484 is in the same RBD loop as S477 and is important for antibody interaction, but not for binding to ACE2. The E484K mutation is expected to have a strong effect since it converts a negatively charged into a positively charged residue, which can significantly alter the RBD interaction antibodies. **(H)** N501Y is in the receptor binding motif and interacts with ACE2. The mutation to Y decreases the hydrophilicity and will enable the formation of pi-stacking interaction with ACE2 residues. **(I)** S494 sits at the tip of a beta-strand in the receptor binding motif. Its mutation to P will likely destabilize the secondary structure.

**Fig. S5.**
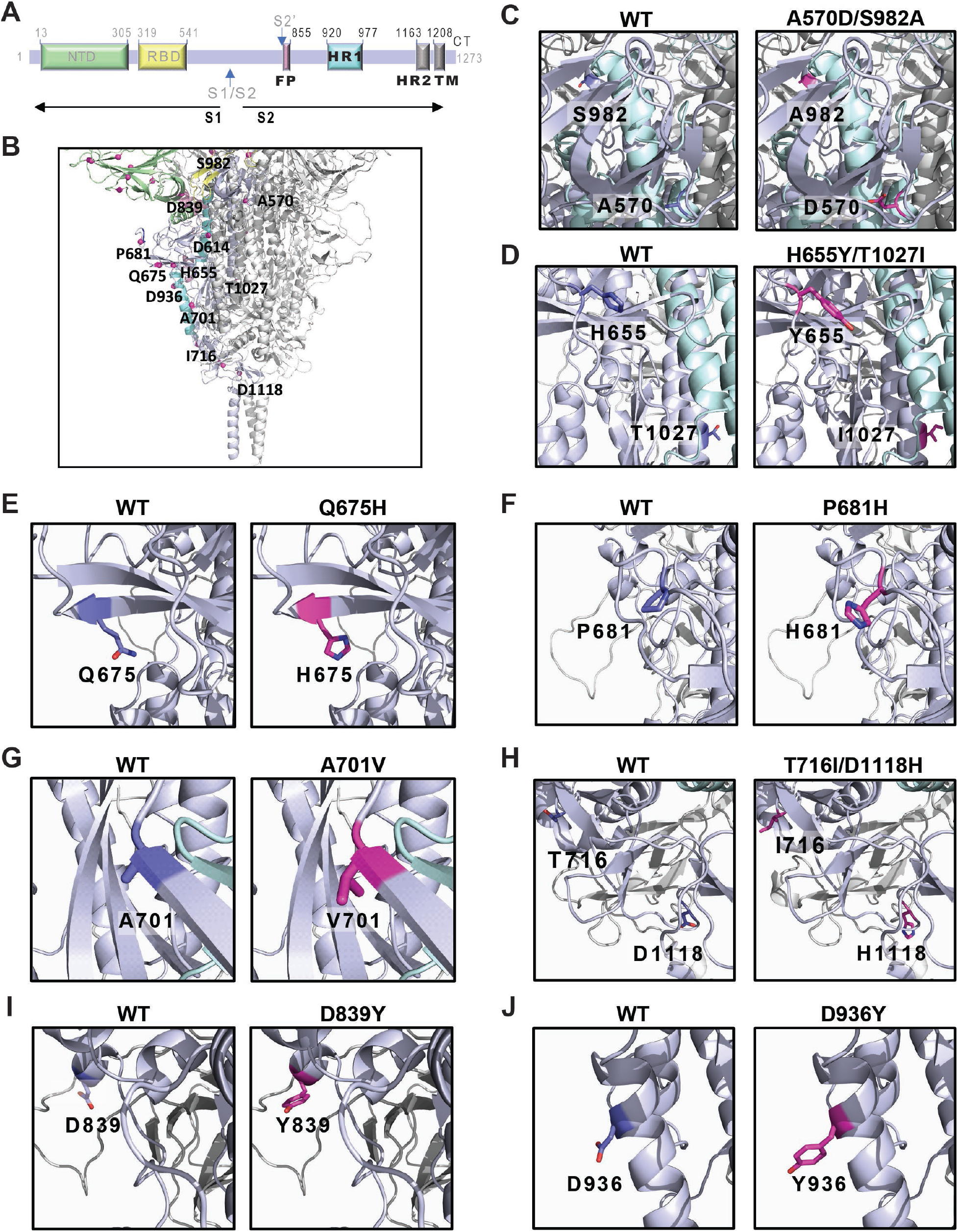
Mutations in the S2 domain of spike. **(A)** Schematic representation of the SARS-CoV-2 spike primary structure. **(B)** The mutations in spike S2 engineered in this study are mapped onto the spike protein surface. Mutations are highlighted using magenta spheres. **(C)** A570 is in an interior loop, the mutation A570D may destabilize this region since the aspartate will face the hydrophobic core of the protein. S982 is located at the tip of an interior alpha-helix and the mutation to A is not predicted to have a strong effect. **(D)** H655 is part of an exposed loop and interacts with F641 through pi-stacking, which can be maintained upon the mutation to Y. T1027 is in an internal alpha-helix and its mutation to I is not predicted to have a significant impact. **(E)** Q675 is present in a beta-sheet and the mutation to H is not predicted to have a significant effect. **(F)** Residue P681 sits in an exposed loop, near the S1 cleavage site. The mutation to H may make this loop more flexible and affect cleavage by proteases and potentially also antibody binding. **(G)** A701 is in an exposed beta-sheet and the mutation to V may destabilize the secondary structure. **(H)** T716 is in an exposed loop and forms a hydrogen bond with the main chain of Q1045, which is lost upon mutation to I. D1118 is found in an interior loop, with the three copies of this residue (one in each monomer) facing one another and this motif can be maintained with the mutation to H. **(I)** Residue D839 sits in an exposed helix turn, very near the fusion peptide, in a region that may be a good target for antibodies. Its mutation to Y is not expected to change the protein structure but may impact the fusion peptide interaction with the host membrane and/or antibody neutralization. **(J)** D936 is in an exposed alpha-helix. The mutation to Y is not expected to change the protein structure but may impact the interaction with antibodies that may bind here.

**Fig. S6.**
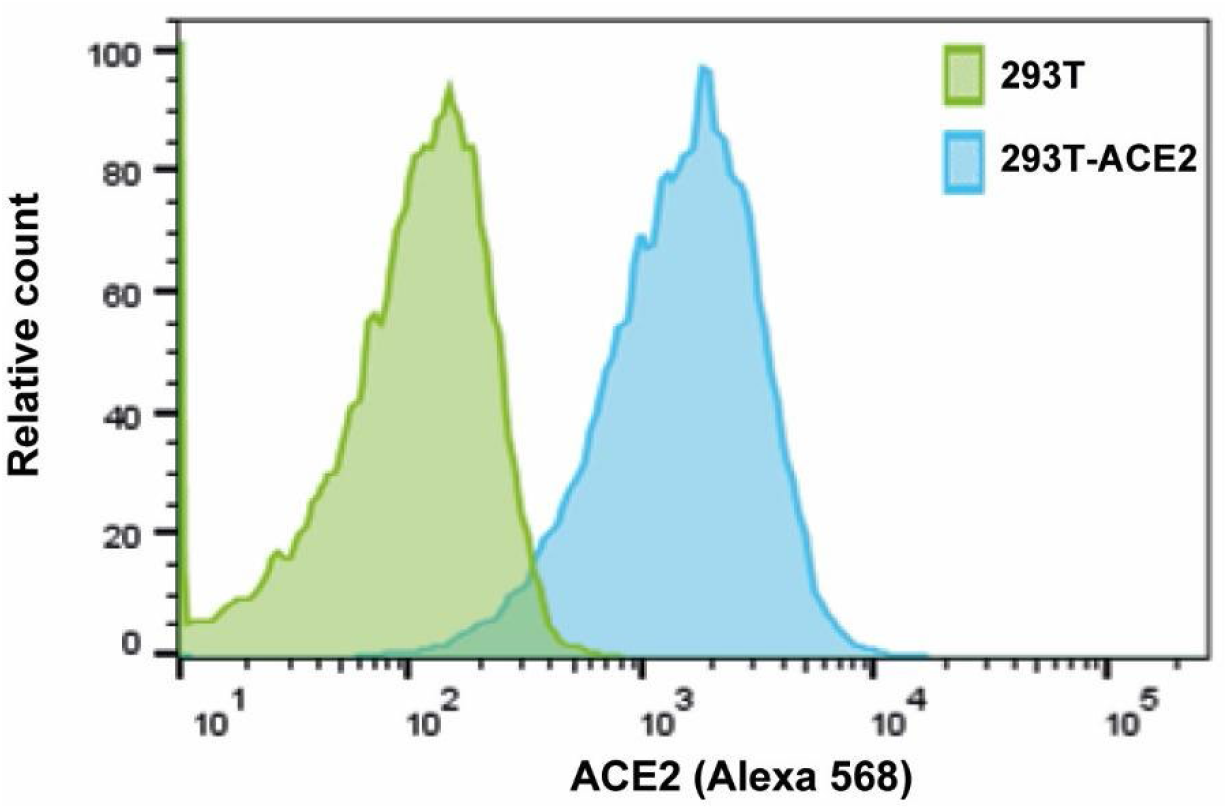
Histogram showing expression of human ACE2 by 293T-ACE2 cell line (blue) compared to parental 293T cells (green). ACE2 expression was measured by flow cytometry, after staining with a goat anti-ACE2 antibody (R&D Systems) followed by staining with an anti-goat antibody conjugated to Alexa 568 (Life Technologies).

**Fig. S7.**
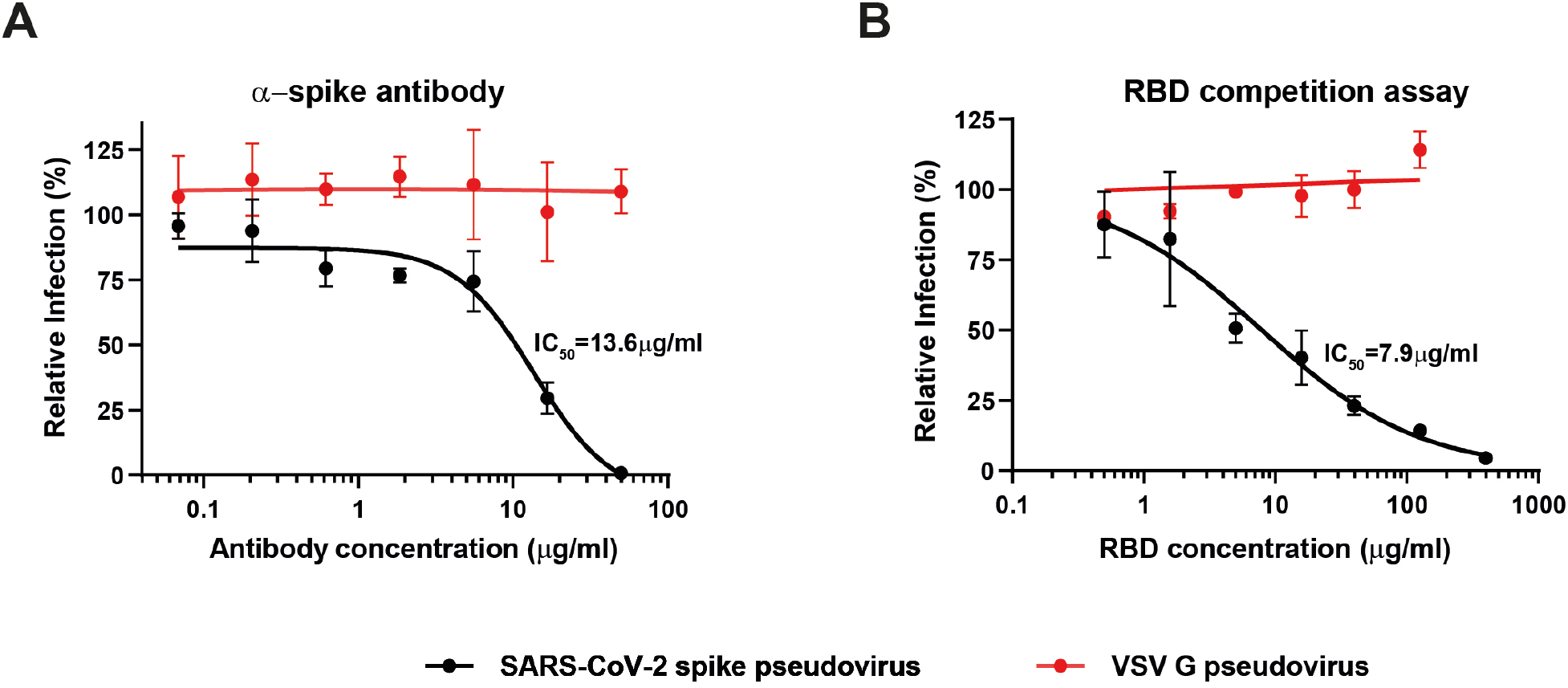
Specificity of the *in vitro* neutralization assay with SARS-CoV-2 spike pseudotyped lentiviral particles. **(A)** SARS-CoV-2 spike specific antibody was tested for neutralization activity against SARS-CoV-2 spike and vesicular stomatitis virus (VSV) G pseudotyped lentivirus. **(B)** Neutralization assay of SARS-CoV-2 spike and VSV G pseudoviruses in the presence of spike’s receptor binding domain (RBD). 293T-ACE2 cells were pre-incubated with serial dilutions of RBD before infection with each virus. Half maximal inhibitory concentration (IC_50_) was calculated for both assays, when possible.

**Fig. S8.**
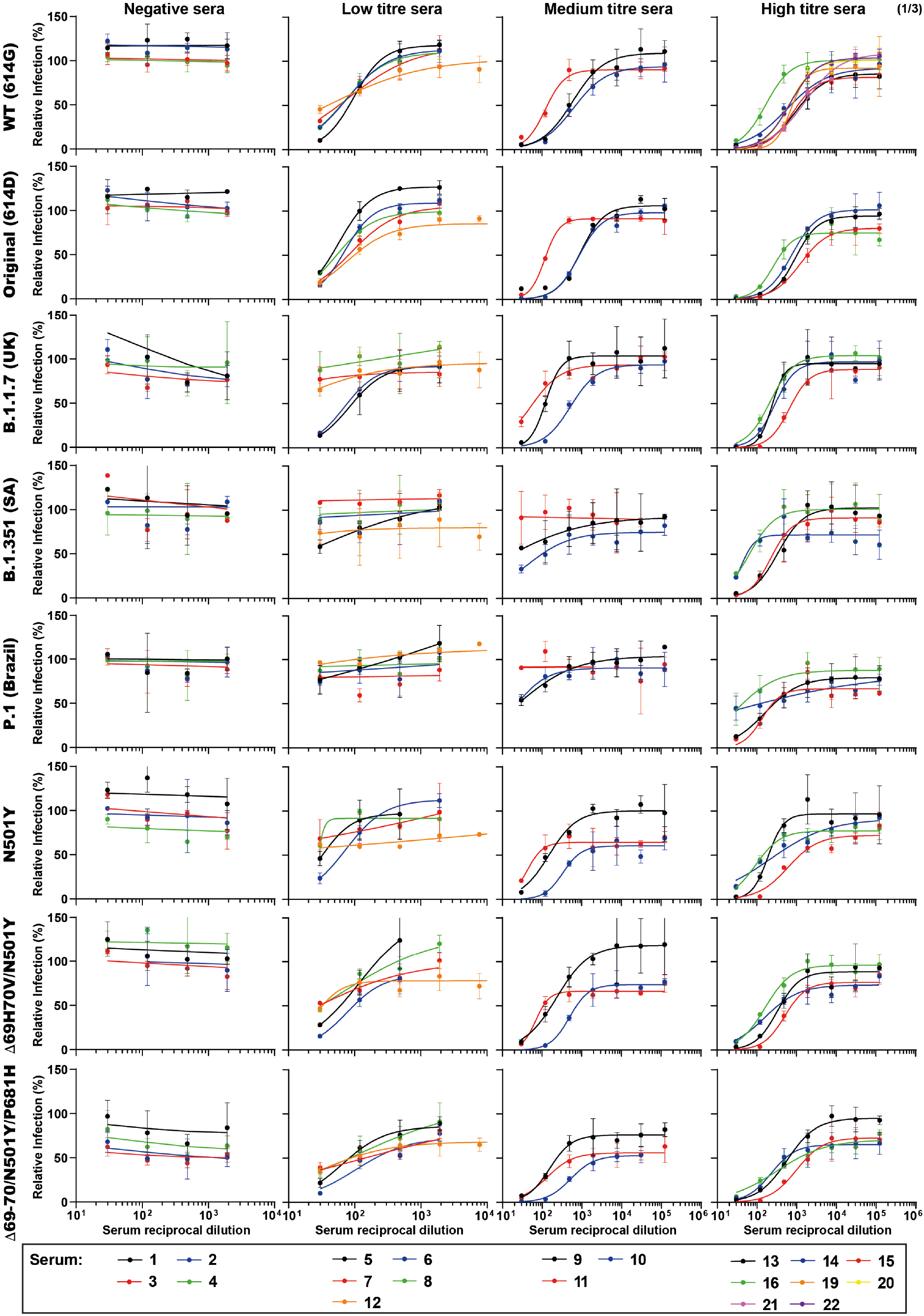
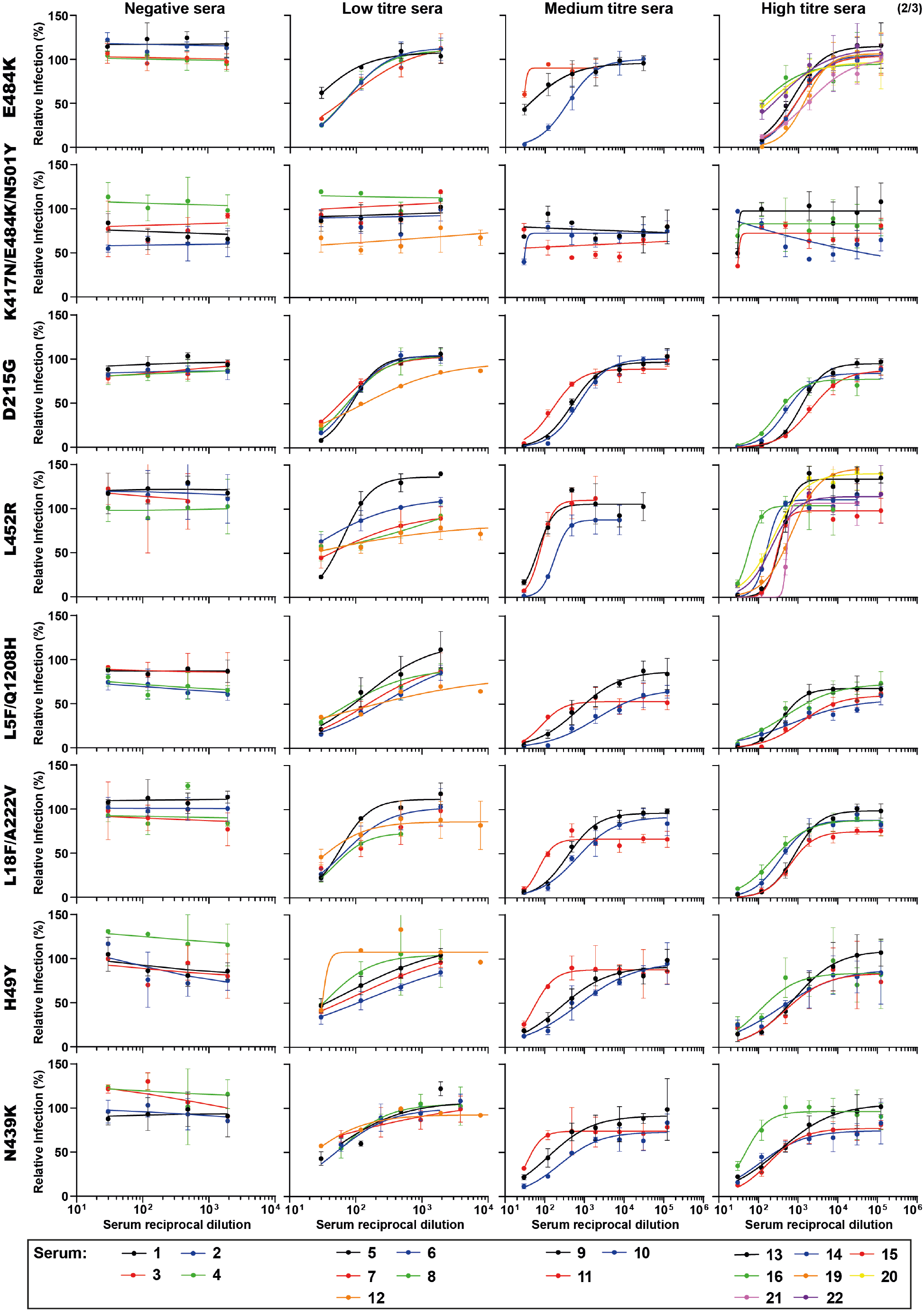
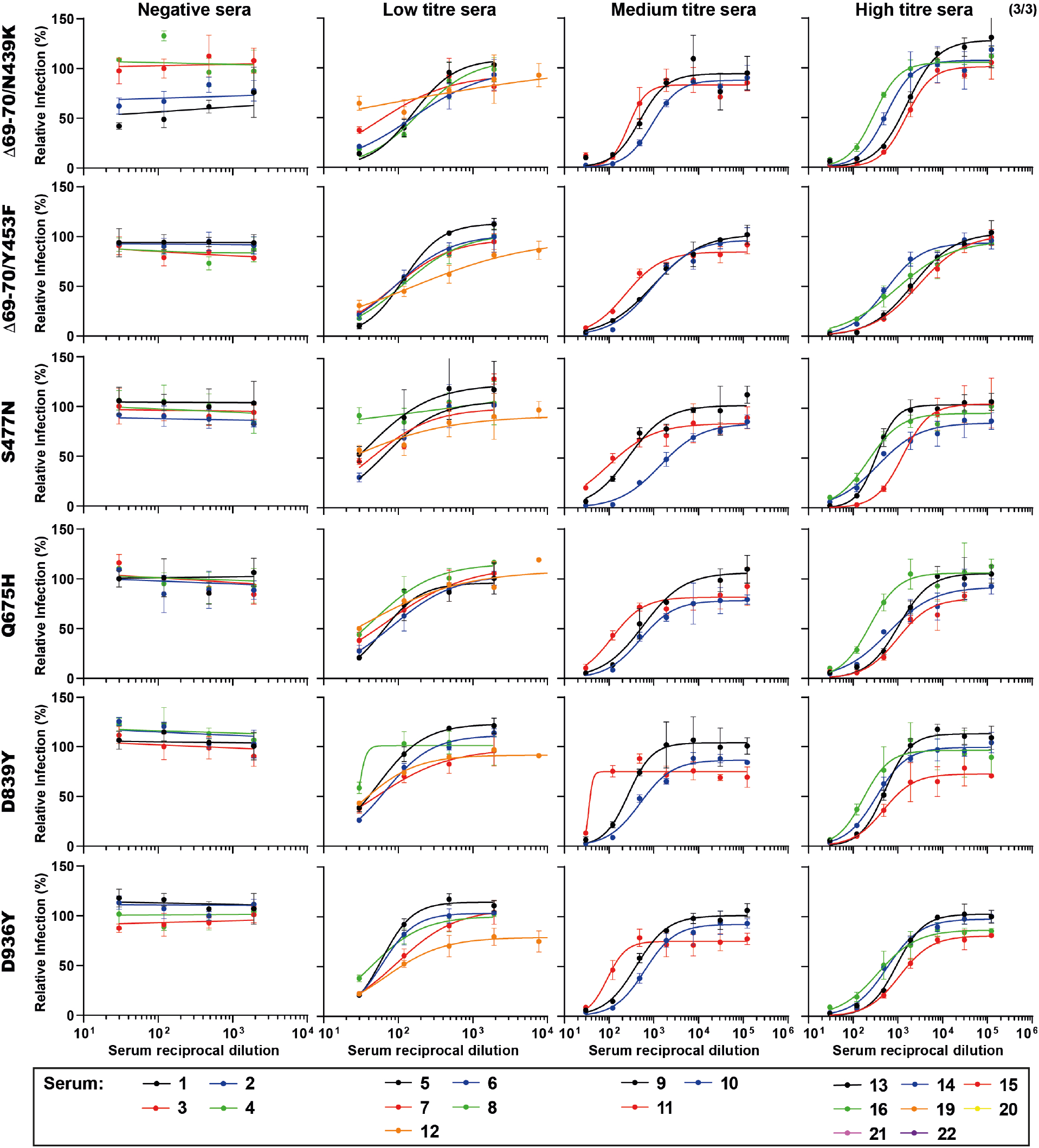
**Neutralization curves of spike mutants by human sera. Related to Fig. 2, Table S2 and Fig. S9.** Sera from 16-20 individuals were tested for neutralization of WT and 22 viruses. Sera were classified into 4 categories: Negative, Low anti-spike IgG titer (≤1:150), Medium titer (1:450) and High titer (≥1:1350). Triplicates were performed for each tested serum dilution. Error bars represent standard deviation.

**Fig. S9.**
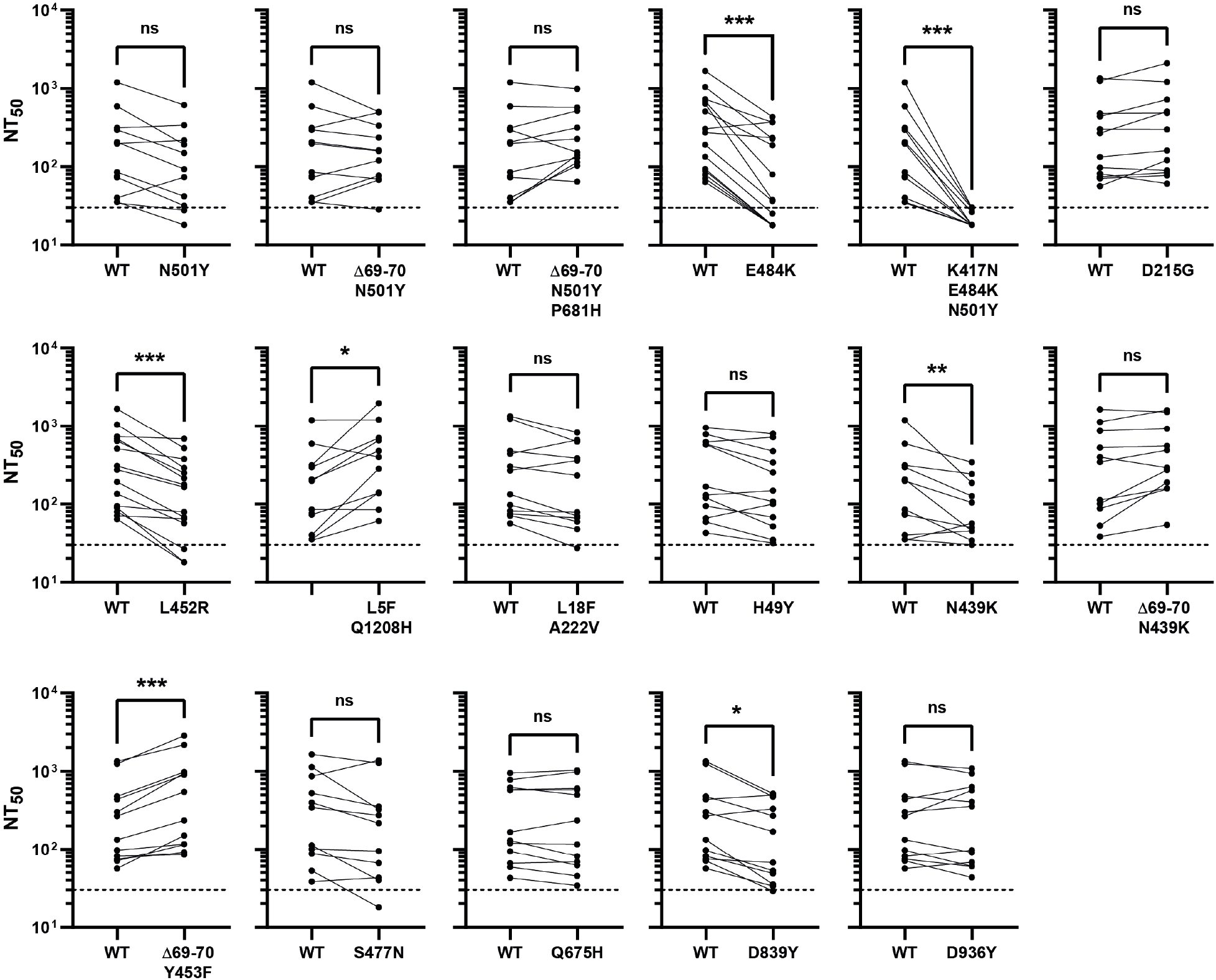
**Neutralization titers of human sera against spike mutants. Related to Fig. 2, Table S2 and fig. S8.** Paired analysis of neutralizing activity of each convalescent serum against WT vs mutant virus. NT_50_ is defined as the inverse of the dilution that achieved a 50% reduction in infection. Dashed lines indicate the limit of detection of the assay (NT_50_=30). ns, non-significant, *p<0.05, **p<0.01, ***p<0.001 by two-tailed Wilcoxon matched-pairs signed-rank test.

**Fig. S10.**
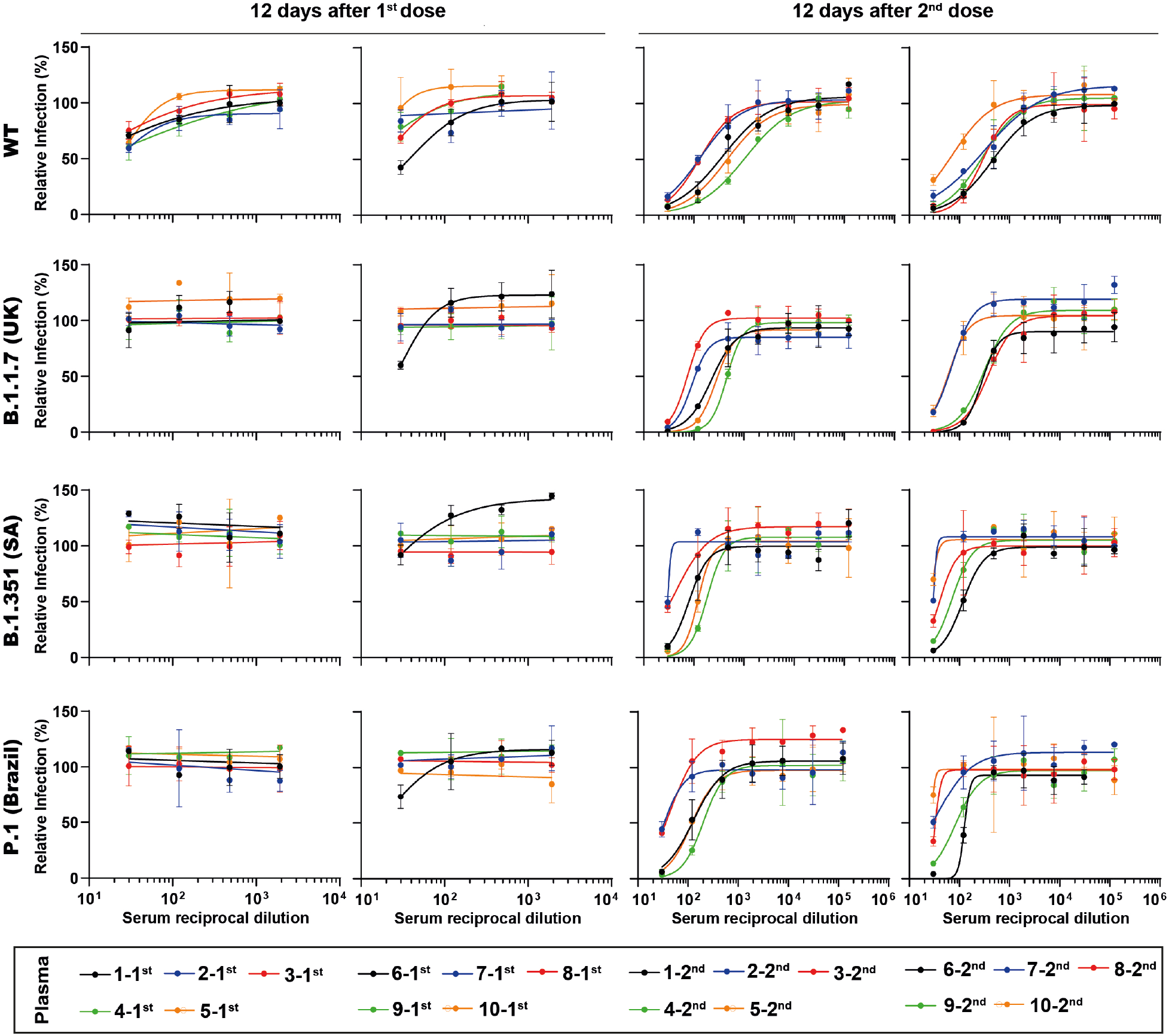
**Neutralization curves of post-vaccination plasma against spike mutants. Related to Fig. 3 and Table S3.** Plasma was collected from 10 individuals 12 days after the first and the second rounds of vaccination and was tested for neutralization of WT virus and variants of concern. Triplicates were performed for each tested plasma dilution. Error bars represent standard deviation.

**Fig. S11.**
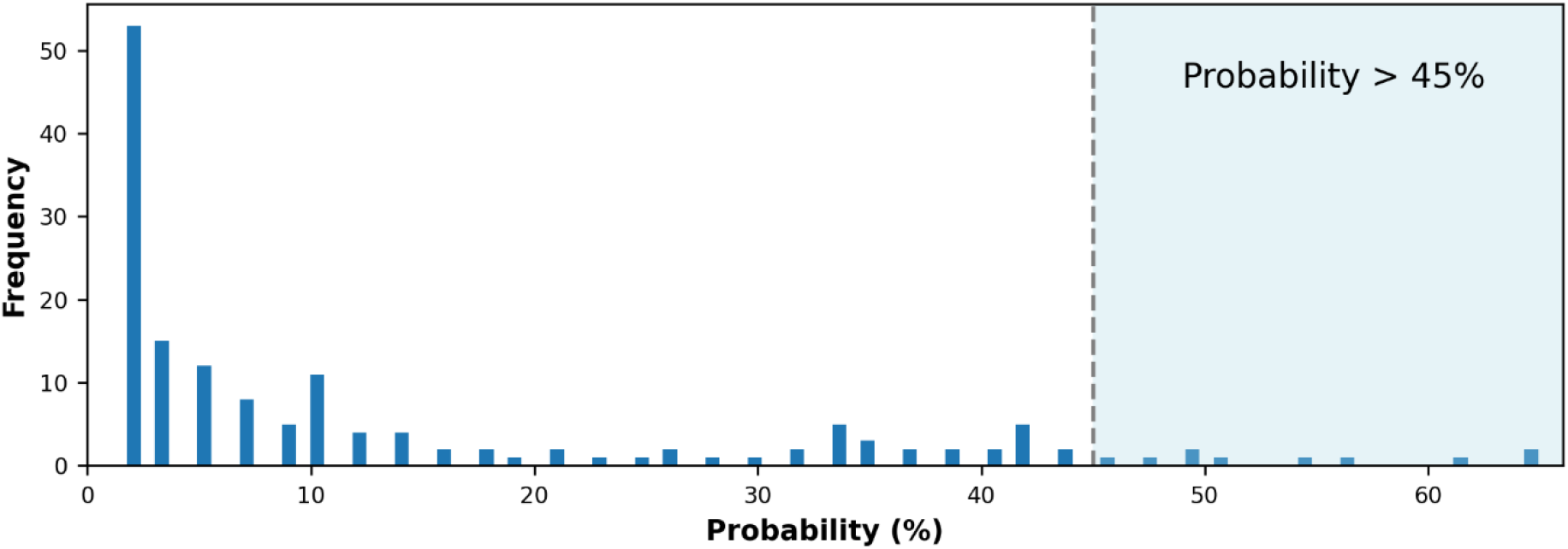
S protein-antibody contact frequency histogram. This was computed by dividing the frequency of interaction of S protein residues with antibodies in 100 bins and calculating the fraction of residues that fall into each bin.

**Figure S12.**
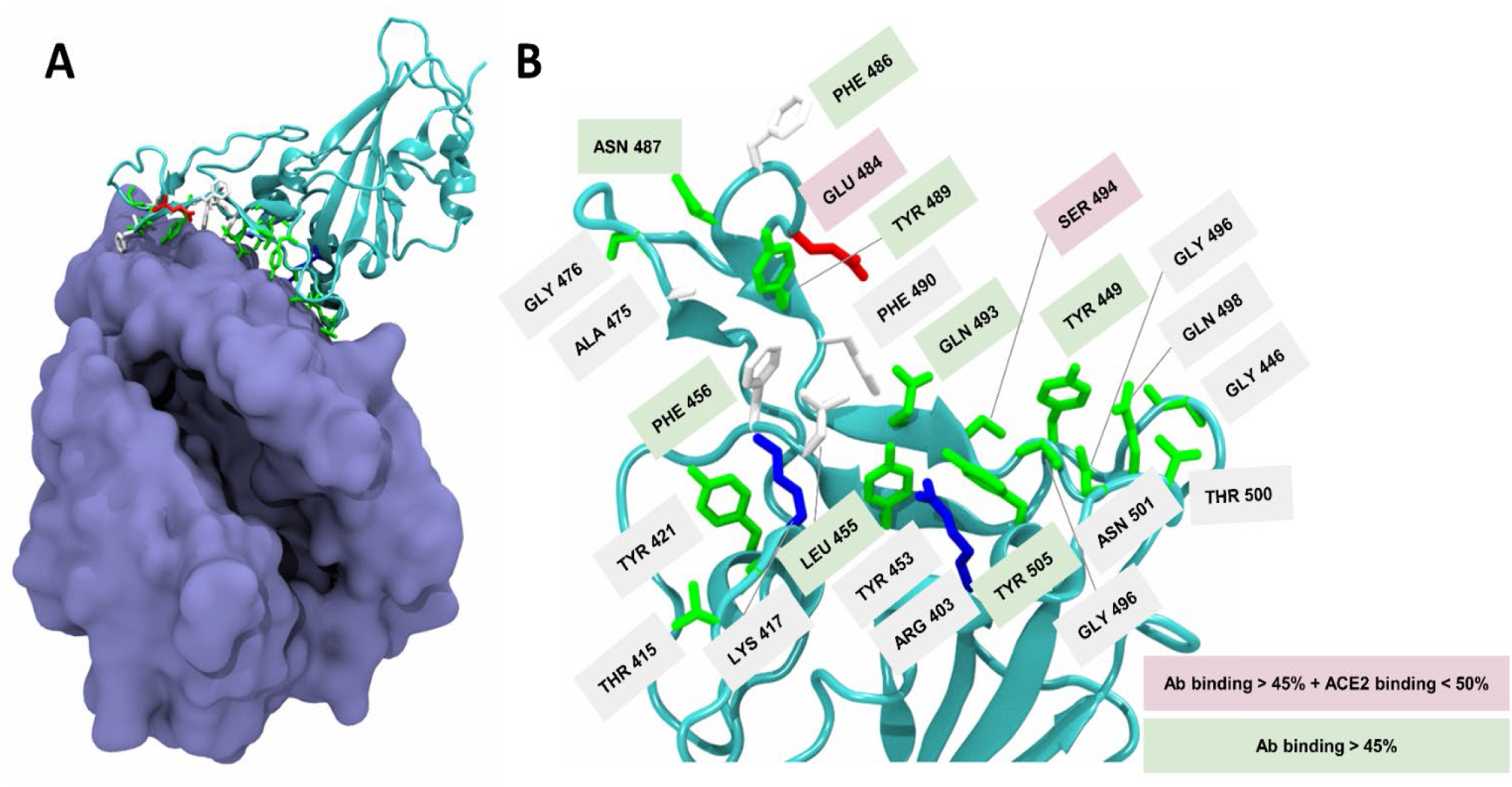
Relevant residues for antibody and ACE2 binding. **(A)** Crystal structure of the SARS-CoV-2 RBD complexed with ACE2. ACE2 is shown as a purple surface and the RBD is shown in a cyan cartoon representation with key antibody interacting residues depicted as sticks. **(B)** Zoom in to the RBM region of the RBD. Residues relevant for antibody binding (>35% frequency of contact) are depicted as sticks. Of these, the ones with an antibody binding probability higher than 45% have a green label, and those that also have a low frequency of binding to ACE2 (<50%) are labelled in pink.

**Fig. S13.**
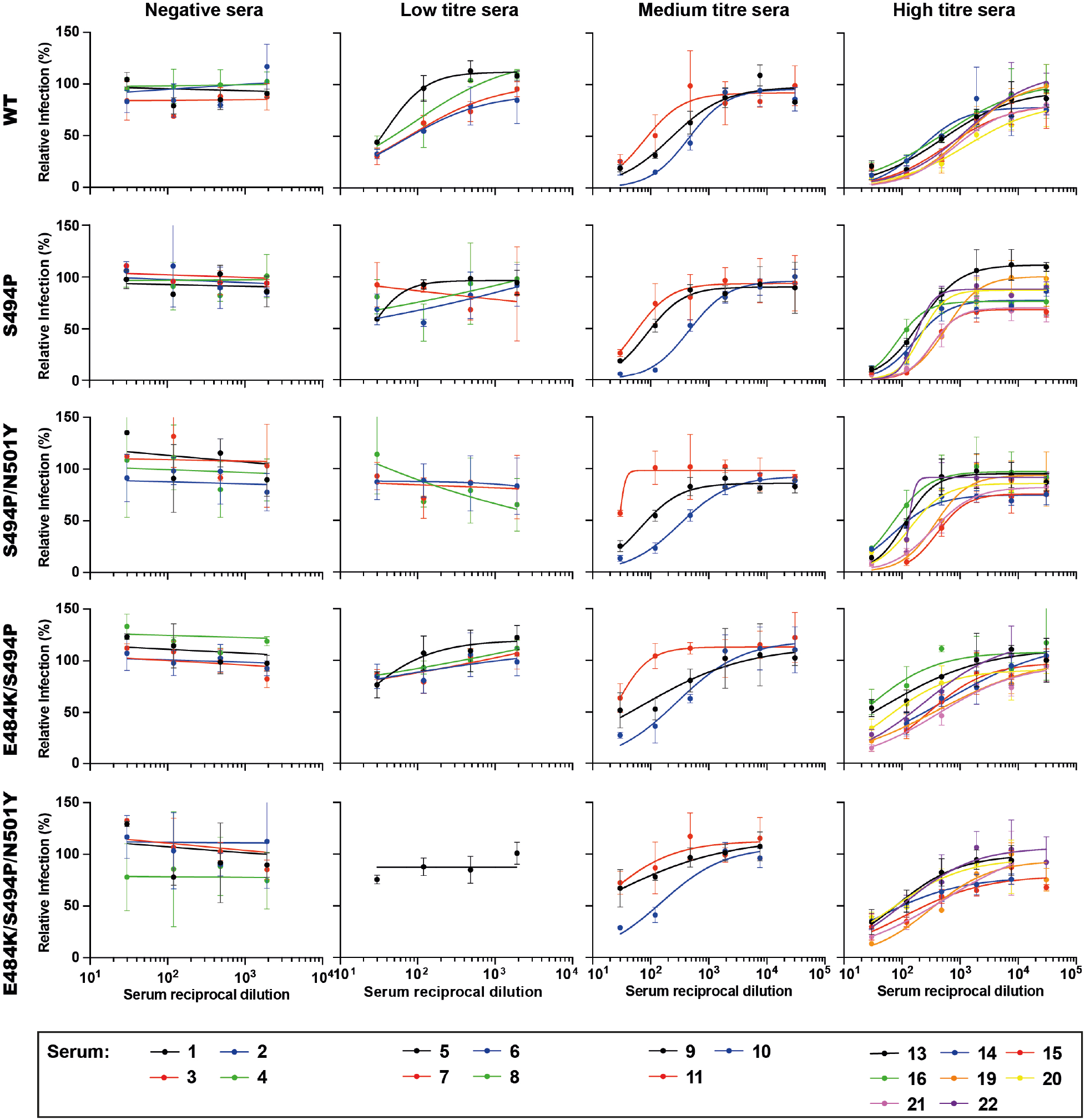
**Neutralization curves of WT and S494P spike mutants by human sera. Related to Fig. 4 and Table S5.** Sera from 19 individuals were tested for neutralization of WT and S494P mutant viruses. Sera were classified into 4 categories: Negative, Low anti-spike IgG titer (≤1:150), Medium titer (1:450) and High titer (≥1:1350). Triplicates were performed for each tested serum dilution. Error bars represent standard deviation.

**Fig. S14.**
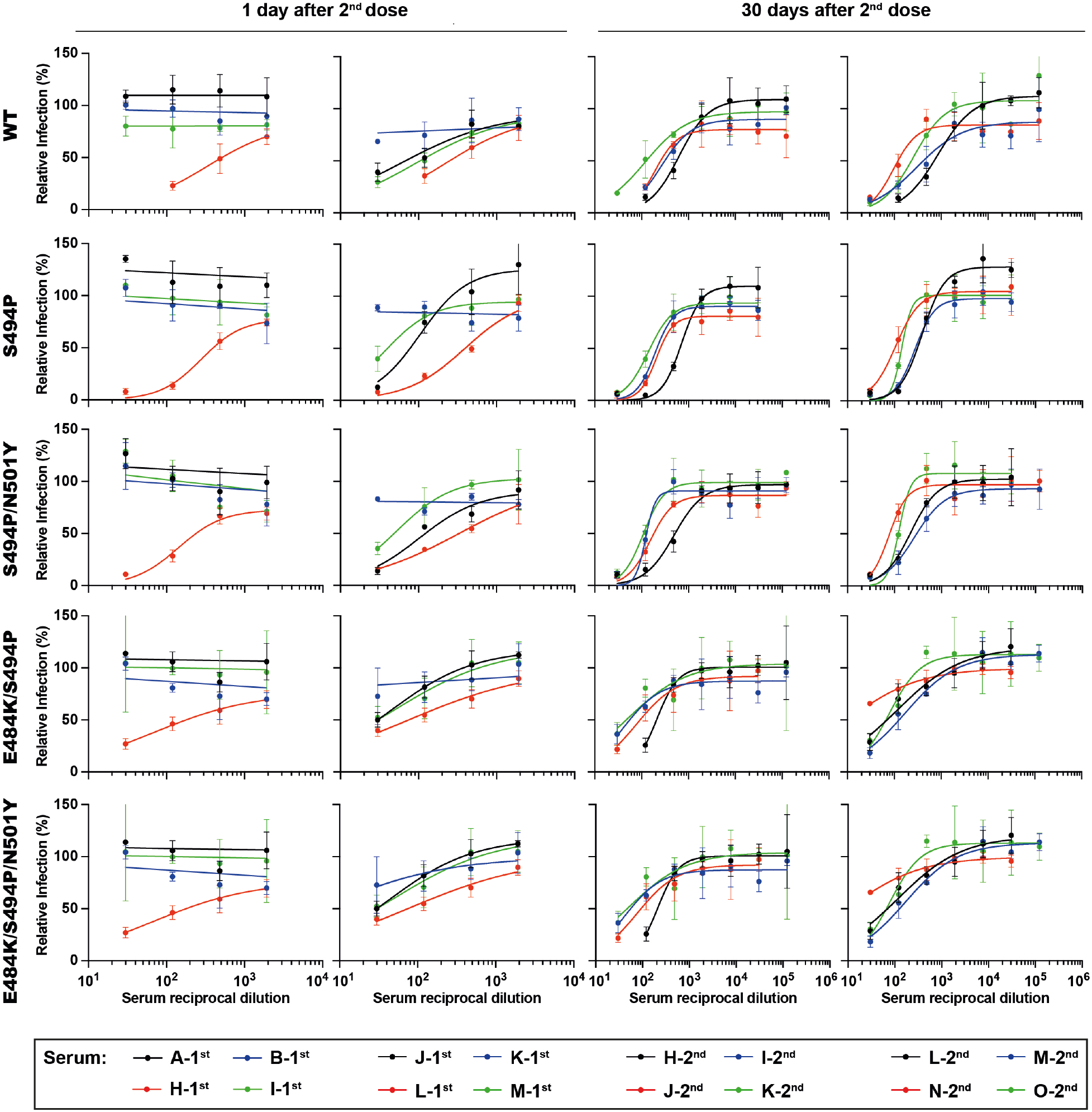
**Neutralization curves of post-vaccination serum against spike mutants. Related to Fig. 4 and Table S6.** Serum was collected from 8 individuals 1 and 30 days after the second round of vaccination, and was tested for neutralization of WT virus and S494P mutants. Triplicates were performed for each tested serum dilution. Error bars represent standard deviation.

**Supplementary Table S1.** Details of the mutants created and used in this study. Values of current frequency are relative to available sequences of week 11 of 2021 (with a bias corresponding to the efforts that countries/regions deploy to SARS-CoV-2 genome surveillance).

| # | Amino acid mutations | References/description |
| --- | --- | --- |
| 1 | D614G<br>From the original to the WT | This mutation emerged early in the epidemic and spread rapidly through Europe and North America, becoming dominant. This is the new WT, and all the other mutations are made on this genotype. Several lines of evidence now suggest that SARS-CoV-2 variants carrying this mutation have increased viral entry, infectivity and transmissibility but not different pathogenicity and do not escape antibody mediated immunity <sup>21,22,109</sup>.<br>Structural information: D614 Sits in a loop. The mutation to Glycine destroys a salt bridge with Lys 864. It will likely make the loop more mobile. There is a glycan nearby.<br>FREQUENCY 98.4 % |
| 2 | Δ69H-70V/Δ144/<br>N501Y/A570D/<br>P681H/T716I/S982A/<br>D1118H | Lineage B.1.1.7 (20I/501Y.V1 (UK)) – all defining mutations.<br>FREQUENCY 65.5% |
| 3 | D80A/D215G/K417/<br>E484K/N501Y/A701V | Lineage B.1.351/501Y.V2 (South Africa) - all defining mutations.<br>FREQUENCY 1.9% |
| 4 | L18F/T20N_P26S/<br>D138Y/R190S/K417T/<br>E484K/N501Y/H655Y/<br>T1027I | Lineage P.1/501Y.V3 (Brazil) - all defining mutations. This lineage was first detected in Manaus in December 2020.<br>FREQUENCY 0.47% |
| 5 | N501Y | Feature mutation in lineages B.1.1.7 (20I/501Y.V1 (UK)); B.1.351/501Y.V2; P.1/501Y.V3. Amino acid 501 is located in the RBD. The N501Y mutation stabilizes virus-binding residues on ACE2 <sup>52,110</sup>. There is convincing evidence that N501Y increases viral transmission <sup>111</sup>.<br>FREQUENCY 71.4% |
| 6 | Δ69H-70V/N501Y | Found in lineage B.1.1.7 (20I/501Y.V1 (UK)) and are considered its most relevant mutations. N501Y was described above. The deletion 69-70 (ΔH69/V70) causes spike gene target failure (SGTF) in some RT-PCR assays <sup>112</sup> and was reported to enhance viral infectivity <sup>43</sup>. Importantly, ΔH69/V70 frequently associates with the mutations N501Y, N439K and Y453F at the RBD, and is also frequently found in association with mutation P681H <sup>43</sup>.<br>FREQUENCY 65.9% |
| 7 | Δ69H-70V/N501Y/<br>P681H | Found in lineage B.1.1.7 and are considered the principal spike mutations. Has spread to multiple countries and its frequency is increasing. Mutations Δ69H-70V and N501Y were described before. P681H is present in Lineage B.1.1.7 and other circulating variants. It was reported as being able to reduce antibody recognition <sup>59</sup>. In addition, it is localized in an exposed loop (677-688) of significant structural flexibility that is unmodelled in several published structures <sup>43</sup>.<br>FREQUENCY 65.9% |
| 8 | E484K | E484 is in the RBD/RBM and interacts with ACE2 at lysine 31, although with low frequency. It has been associated with antibody escape from therapeutic antibodies, natural infection, convalescent sera and vaccination-induced immunity <sup>36,57,83</sup>.<br>FREQUENCY 6.4% |
| 9 | K417N/E484K/N501Y | Mutation present in lineages B.1.351/501Y.V2 (South Africa) (lacks mutations D80A, D215G and A701V) and P.1/501Y.V3 (Brazil) that, in most cases, instead of K417N has the mutation K417T. However, as T and N are similar amino acids, the mutations have similar effects. Deep mutational scanning indicates that N or T change to K in position 417 mutation has minimal effect on spike binding affinity to ACE2 <sup>4</sup>. K417 is a unique ACE2 interacting residue that forms a salt bridge interaction across the central contact region with D30 of ACE2 <sup>110</sup>.<br>FREQUENCY 2.0% |
| 10 | L5F/Q1208H | L5F - Mutations in the N terminal have been shown to affect neutralization <sup>54</sup>.<br>There is no structural information for any of these amino acids because they localize to unstructured regions at the NTD, TD and CD and therefore it is not possible to map them in fig S3-5.<br>FREQUENCY 0.0% |
| 11 | L18F/A222V | L18F – Present in lineage. B.1.1.28.1/P.1. Has a mild impact in the structure of spike and also protects from some neutralizing monoclonal antibodies <sup>54</sup>. A222V – Mutation that expanded in Europe in lineage B.1.117 and is not associated with antibody escape <sup>54,113</sup>. These mutations are frequently found in association.<br>FREQUENCY 0.9% |
| 12 | H49Y | Mutation identified in China <sup>114</sup> and not reported to affect the function of spike <sup>109</sup>.<br>FREQUENCY 0.3% |
| 13 | D215G | This mutation is present in Lineage B.1.351 and is one of many mutations in the NTD of spike reported to contribute to resistance to neutralizing antibodies <sup>54,115</sup>.<br>This mutation is currently found exclusively associated with B. 1.351. However, its prevalence in the past has displayed two peaks unrelated to E484K (see prevalence in fig. S2).<br>FREQUENCY 2.2% |
| 14 | N439K | Found in lineages B.1.141 (common in UK in the beginning of pandemics, until June) and B.1.258 (appeared in April, and its presence has been slowly increasing). N439K mutation sits in the RBD and results show that the virus retains viral fitness but becomes resistant to some neutralizing antibodies <sup>41,116</sup>. The N439 sits at the extremity of RBD, but not directly in the contact zone to ACE2. It is under the residues at the contact zone. It is exposed in the "up" conformation of the RBD. In the down conformation it is semi-exposed. A likely zone for antibody binding.<br>FREQUENCY 2.0% |
| 15 | Δ69H-70V/N439K | Δ69-70 and N439K have been explained individually above. Importantly, ΔH69/V70 frequently associates with the mutations in RBD N501Y, N439K and Y453F <sup>43</sup>.<br>FREQUENCY 2.0% |
| 16 | L452R | Part of lineage B.1.427/9 / 20C/S (USA CAL). It is located in RBD and has been shown to increase infectivity <sup>57</sup> and escape antibody binding in a screen in yeast using the RBD and not full-length spike <sup>2</sup> and in <sup>60</sup>.<br>FREQUENCY 4.2% |
| 17 | Δ69H-70V/Y453F | These set of mutations were found together in association with mink related infections. It was found initially in Denmark and the Netherlands and resulted in culling of minks <sup>40</sup>. Δ69-70 was explained above. Y453 is localised in the RBD and was shown to affect neutralization by SARS-CoV-2 specific antibodies <sup>116</sup>. It requires better understanding.<br>FREQUENCY 0.0% |
| 18 | S477N | Localized at the RBD, was reported to have a modest increase in the affinity of the RBD to ACE2 <sup>4</sup>.<br>FREQUENCY 2.4% |
| 19 | Q675H | Q675H leads to a putative change in glycosylation and was reported to reduce infectivity <sup>57</sup>.<br>FREQUENCY 0.20% |
| 20 | D839Y | Prevalent in Portugal by 30<sup>th</sup> April <sup>117</sup>. The aspartic acid 839 is exposed, not making relevant interactions. It may be a zone for antibody targeting. This zone may be important for fusion and/or in the induction of host inflammatory responses.<br>FREQUENCY 0.01% |
| 21 | D936Y | The aspartic acid 936Y is located in the fusion core of the heptad repeat 1. It was detected in Sweden and England and was reported to destabilize the post-fusion conformation of spike, while minimally impacting on the stability of the pre-fusion <sup>118</sup>. A distinct paper reported that this mutation increased infectivity but did not affect neutralization by antibodies <sup>119</sup>.<br>The aspartic acid 936 sits in an exposed helix. Its substitution for a tyrosine looks harmless from the perspective of structural stability.<br>FREQUENCY 0.7% |
| 22 | S494P | The acquisition of Spike mutation S494P in was observed for B.1.1.7 at the end of February 2021, and was acquired multiple times, and its frequency is increasing. S494P allows evasion of binding or neutralization by several monoclonal antibodies <sup>120</sup> but has not been shown to impact on neutralization by convalescent or vaccine-induced polyclonal antisera. S494P confers increased binding to hACE2 <sup>4</sup>.<br>FREQUENCY 0.81% |
| 23 | S494P/N501Y | The acquisition of Spike mutation S494P in was observed for B.1.1.7 at the end of February 2021, and was acquired multiple times, and its frequency is increasing<br>FREQUENCY 0.44% |
| 24 | E484K/S494P | Although still at a very low frequency, this double mutation is starting to appear.<br>FREQUENCY 0.01% |
| 25 | E484K/S494P/N501Y | This triple mutation was not detected yet.<br>FREQUENCY 0.00% |

**Table S2.** IgG antibody titers against SARS-CoV-2 spike protein and neutralizing titers (NT_50_) of convalescent sera against WT and mutant spike pseudoviruses.

| Serum ID | ELISA IgG titre |  | NT<sub>50</sub> (95% confidence interval) |  |  |  |  |  |  |  |  |  |  |  |
| --- | --- | --- | --- | --- | --- | --- | --- | --- | --- | --- | --- | --- | --- | --- |
|  |  |  | WT (614G) | Original (614D) | B.1.1.7 (U.K.) | B.1.351 (S.A.) | P.1 (Brazil) | N501Y | Δ69-70 N501Y | Δ69-70/N501Y P681H | E484K | K417N/E484K N501Y | L452R | S494P |
| 1* | Negative control | <50 | <30 | <30 | <30 | <30 | <30 | <30 | <30 | <30 | <30 | <30 | <30 | <30 |
| 2 |  | <50 | <30 | <30 | <30 | <30 | <30 | <30 | <30 | <30 | <30 | <30 | <30 | <30 |
| 3 |  | <50 | <30 | <30 | <30 | <30 | <30 | <30 | <30 | <30 | <30 | <30 | <30 | <30 |
| 4 |  | <50 | <30 | <30 | <30 | <30 | <30 | <30 | <30 | <30 | <30 | <30 | <30 | <30 |
| 5 | Low titre | 50 | 97 (83-114) | 57 (48-68) | 84 (41-545) | <30 | <30 | 32 (?-?) | 120 (34-?) | 64 (37-304) | <30 | <30 | 65 (52-80) | <30 |
| 6 |  | 150 | 74 (51-119) | 71 (61-82) | 70 (48-102) | <30 | <30 | 73 (55-99) | 77 (41-?) | 103 (42-?) | <30 | <30 | <30 | <30 |
| 7 |  | 150 | 81 (41-128) | 87 (59-184) | <30 | <30 | <30 | 30 (?-?) | 28 (13-46) | 114 (16-?) | <30 | <30 | <30 | <30 |
| 8 |  | 150 | 71 (53-102) | 53 (42-68) | <30 | <30 | <30 | 28 (0-?) | 68 (25-?) | 141 (60-283) | <30 | <30 | <30 | <30 |
| 12 |  | 150 | 43 (19-94) | 76 (55-105) | <30 | <30 | <30 | <30 | 25 (1-36) | 26 (5-115) | n.d. | <30 | <30 | <30 |
| 9 | Medium titre | 450 | 573 (242-3770) | 924 (696-1250) | 123 (75-204) | <30 | 28 (7-167) | 149 (75-310) | 235 (118-522) | 153 (101-236) | 37 (14-115) | <30 | 68 (44-?) | 55 (39-77) |
| 10 |  | 450 | 621 (407-1019) | 854 (677-1105) | 527 (396-720) | 41 (10-556) | 19 (2—35) | 339 (166-744) | 492 (357-696) | 520 (365-797) | 372 (264-545) | 29 (21-33) | 177 (?-258) | 154 (122-199) |
| 11 |  | 450 | 120 (92-152) | 117 (101-135) | 53 (33-95) | <30 | <30 | 42 (26-69) | 69 (51-?) | 130 (74-254) | <30 | <30 | 79 (47-?) | 30 (28-35) |
| 13 | High titre | 4050 | 954 (574-1606) | 946 (762-1180) | 241 (?-?) | 339 (183-589) | 161 (80-345) | 192 (122-340) | 333 (204-536) | 572 (430-776) | 188 (126-312) | 30 (?-?) | 376 (308-?) | 293 (213-408) |
| 14 |  | 4050 | 581 (278-1674) | 769 (576-1052) | 283 (170-454) | 40 (24-70) | 27 (1-?) | 217 (107-761) | 159 (95-312) | 226 (127-440) | 236 (159-388) | <30 | 165 (?-201) | 131 (87-209) |
| 15 |  | 4050 | 783 (655-944) | 1248 (855-1859) | 663 (471-1006) | 226 (136-388) | 137 (80-218) | 613 (351-1579) | 504 (374-715) | 984 (590-1660) | 229 (166-330) | 30 (24-37) | 288 (204-?) | 397 (281-597) |
| 16 |  | 1350 | 167 (120-235) | 263 (159-433) | 217 (167-281) | 66 (31-125) | 32 (2-1.1E6) | 93 (62-141) | 162 (118-223) | 315 159-1052) | <30 | <30 | 57 | 60 (40-89) |
| 19 |  | 4050 | 729 (?-1075) | n.d. | n.d. | n.d. | n.d. | n.d. | n.d. | n.d. | 371 (245-553) | n.d. | 686 (503-982) | 345 (259-482) |
| 20 |  | 1350 | 641 (528-789) | n.d. | n.d. | n.d. | n.d. | n.d. | n.d. | n.d. | 38 (6-2.9E7) | n.d. | 250 (177-355) | 283 (209-390) |
| 21 |  | 1350 | 1668 (124-2333) | n.d. | n.d. | n.d. | n.d. | n.d. | n.d. | n.d. | 435 (261-1002) | n.d. | 518 (?-735) | 653 (399-1238) |
| 22 |  | 1350 | 684 (591-797) | n.d. | n.d. | n.d. | n.d. | n.d. | n.d. | n.d. | 80 (26-2727) | n.d. | 213 (144-314) | 261 (173-404) |
| Serum ID | ELISA IgG titre |  | L5F/Q1208H S1252P | L18F A222V | H49Y | D215G | Δ69-70 N439K | N439K | L452R | Δ69-70 Y453F | S477N | Q675H | D839Y | D936Y |
| 1* | Negative control | <50 | <30 | <30 | <30 | <30 | <30 | <30 | <30 | <30 | <30 | <30 | <30 | <30 |
| 2 |  | <50 | <30 | <30 | <30 | <30 | <30 | <30 | <30 | <30 | <30 | <30 | <30 | <30 |
| 3 |  | <50 | <30 | <30 | <30 | <30 | <30 | <30 | <30 | <30 | <30 | <30 | <30 | <30 |
| 4 |  | <50 | <30 | <30 | <30 | <30 | <30 | <30 | <30 | <30 | <30 | <30 | <30 | <30 |
| 5 | Low titre | 50 | 138 (47-?) | 59 (46-76) | 68 (25-?) | 90 (77-103) | 158 (114-248) | 50 (10-120) | 60 (58-62) | 117 (106-129) | 40 (5-?) | 62 (39-121) | 53 (44-64) | 60 (51-71) |
| 6 |  | 150 | 282 (118-1.3E5) | 66 (37-351) | 148 (31-?) | 84 (70-99) | 155 (105-225) | 45 (13-?) | <30 | 92 (59-196) | 66 (38-161) | 82 (51-207) | 67 (56-83) | 60 (47-75) |
| 7 |  | 150 | 140 (45-?) | 78 (38-138) | 100 (36-?) | 61 (54-68) | 54 (15-?) | <30 | 37 (2-?) | 85 (66-119) | 43 (12-93) | 69 (33-40536) | 49 (25-7.4E5) | 97 (68-168) |
| 8 |  | 150 | 61 (19-?) | 48 (36-68) | 35 (0-?) | 77 (55-114) | 189 (100-2444) | 56 (5-?) | <30 | 115 (77-237) | <30 | 45 (29-127) | 29 (16-34) | 43 (33-60) |
| 12 |  | 150 | 78 (9-?) | 27 (2-59) | 32 (27-47) | 120 (85-176) | <30 | 20 (4-34) | 168 (124-243) | 150 (98-256) | 17 (1-44) | 34 (7-341) | 34 (17-53) | 68 (44-103) |
| 9 | Medium titre | 450 | 699 (377-1559) | 384 (296-511) | 253 (145-539) | 484 (382-628) | 487 (286-825) | 127 (54-1170) | 307 (213-418) | 982 (698-1474) | 274 (162-590) | 584 (333-1281) | 270 (180-401) | 406 (330-504) |
| 10 |  | 450 | 1973 (820-8651) | 662 (418-1120) | 715 (288-5790) | 721 (561-941) | 926 (749-1149) | 243 (134-608) | 67 (44-?) | 907 (615-1484) | 1379 (1044- | 500 (312-872) | 496 (382-654) | 631 (478-852) |
| 11 |  | 450 | 85 (51-151) | 70 (50-?) | 52 (31-85) | 161 (118-228) | 272 (185-?) | 34 (20-52) | <30 | 235 (171-339) | 95 (54-184) | 116 (72-221) | 35 (?-63) | 91 (65-119) |
| 16 | High titre | 1350 | 477 (199-2315) | 231 (160-344) | 107 (17-5582) | 300 (207-451) | 290 (233-359) | 342 (194-765) | 538 (404-720) | 941 (535-2061) | 216 (161-298) | 236 (149-377) | 170 (99-292) | 936 (786-1116) |
| 13 |  | 4050 | 399 (285-558) | 825 (671-1023) | 793 (421-2152) | 1212 (1057- | 1609 (1212- | 105 (45-1015) | 167 (109-268) | 2170 (1585- | 328 (277-383) | 1032 (738-1453) | 520 (431-628) | 570 (466-706) |
| 14 |  | 4050 | 648 (194-1.8E6) | 358 (271-481) | 339 (81-1.6E7) | 506 (395-662) | 552 (366-878) | 186 (129-275) | 581 (325-1064) | 544 (402-786) | 353 (211-682) | 603 (340-1321) | 331 (262-421) | 1082 (869-1353) |
| 15 |  | 4050 | 1208 (741-2273) | 627 (500-795) | 474 (136-3404) | 2094 (1578- | 1520 (1150- | 45 (27-73) | 135 (68-279) | 2858 (2013- | 1276 (863-1959) | 987 (657-1947) | 475 278-871) | 357 (234-567) |

<sup>a</sup> pre-pandemic pool
? - could not be calculated
n.d. – not done

**Table S3.** IgG antibody titers against SARS-CoV-2 spike protein and neutralizing titers (NT_50_) against WT and variant pseudoviruses of plasma from vaccinated individuals, collected 12 days after the first and after the second doses of the vaccine.

| Plasma ID | ELISA<br>IgG titer | NT<sub>50</sub> (95% confidence interval) |  |  |  |
| --- | --- | --- | --- | --- | --- |
|  |  | WT | B.1.1.7 (UK) | B.1.351 (SA) | P.1 (Brazil) |
| v1-1<sup>st</sup> | 4050 | <30 | <30 | <30 | <30 |
| v2-1<sup>st</sup> | 4050 | 19 (1-31) | <30 | <30 | <30 |
| v3-1<sup>st</sup> | 1350 | <30 | <30 | <30 | <30 |
| v4-1<sup>st</sup> | 4050 | <30 | <30 | <30 | <30 |
| v5-1<sup>st</sup> | 4050 | 25 (22-28) | <30 | <30 | <30 |
| v6-1<sup>st</sup> | 12150 | 40 (24-68) | 31 (21-43) | <30 | <30 |
| v7-1<sup>st</sup> | 4050 | <30 | <30 | <30 | <30 |
| v8-1<sup>st</sup> | 1350 | <30 | <30 | <30 | <30 |
| v9-1<sup>st</sup> | 1350 | <30 | <30 | <30 | <30 |
| v10-1<sup>st</sup> | 450 | <30 | <30 | <30 | <30 |
| v1-2<sup>nd</sup> | 36450 | 380 (210-952) | 224 (158-324) | 79 (52-112) | 131 (87-209) |
| v2-2<sup>nd</sup> | 36450 | 134 (84-222) | 92 (70-?) | 30 (24-38) | 33 (19-49) |
| v3-2<sup>nd</sup> | 109350 | 131 (104-166) | 75 (59-?) | 43 (29-63) | 46 (35-61) |
| v4-2<sup>nd</sup> | 36450 | 1053 (777-1486) | 458 (402-515) | 185 (?-262) | 203 (?-320) |
| v5-2<sup>nd</sup> | 36450 | 457 (288-790) | 277 (227-333) | 122 (92-159) | 117 (90-157) |
| v6-2<sup>nd</sup> | 109350 | 456 (324-663) | 279 (197-?) | 116 (94-143) | 125 (113-164) |
| v7-2<sup>nd</sup> | 36450 | 316 (194-536) | 71 (54-91) | 30 (26-38) | 35 (17-56) |
| v8-2<sup>nd</sup> | 36450 | 290 (189-439) | 381 (285-539) | 40 (26-67) | 33 (?-56) |
| v9-2<sup>nd</sup> | 109350 | 291 (190-460) | 310 (234-409) | 71 (52-95) | 83 (57-120) |
| v10-2<sup>nd</sup> | 136450 | 72 (47-109) | 61 (42-85) | <30 | <30 |

? - could not be calculated

**Supplementary Table S4.** Summary of antibodies studied in the S protein-antibody complexes. These antibodies were chosen due to the availability of their structure resolved together with the S protein (or just the RBD region) in the PDB repository ^94^.

| SARS-CoV-2 antibodies studied |  |  |  |  |  |  |
| --- | --- | --- | --- | --- | --- | --- |
| 15033 | 2-4 | 2-15 | 15033-7 | 2G12 | 2H2 | 3C1 |
| 4A8 | B38 | BD-236 | BD-368-2 | BD-604 | BD-629 | BD23 |
| C002 | C102 | C104 | C105 | C110 | C119 | C121 |
| C135 | C144 | C1A-B12 | C1A-B3 | C1A-C2 | C1A-F10 | CB6 |
| CV30 | DH1047 | DH1050.1 | EY6A | FC05 | H014 | LY-CoV555 |
| CC12.3 | COVA1-16 | COVA2-04 | COVA2-39 | CT-P59 | CV07-250 | CV07-270 |
| P17 | P2B-2F6 | P2C-1A3 | P2C-1F11 | REGN10933 | REGN10987 | S2A4 |
| S2E12 | S2H13 | S2H14 | S2M11 | S304 | S309 | STE90-C11 |
| No name<br>(7cjf) |  |  |  |  |  |  |

**Supplementary Table 5.**
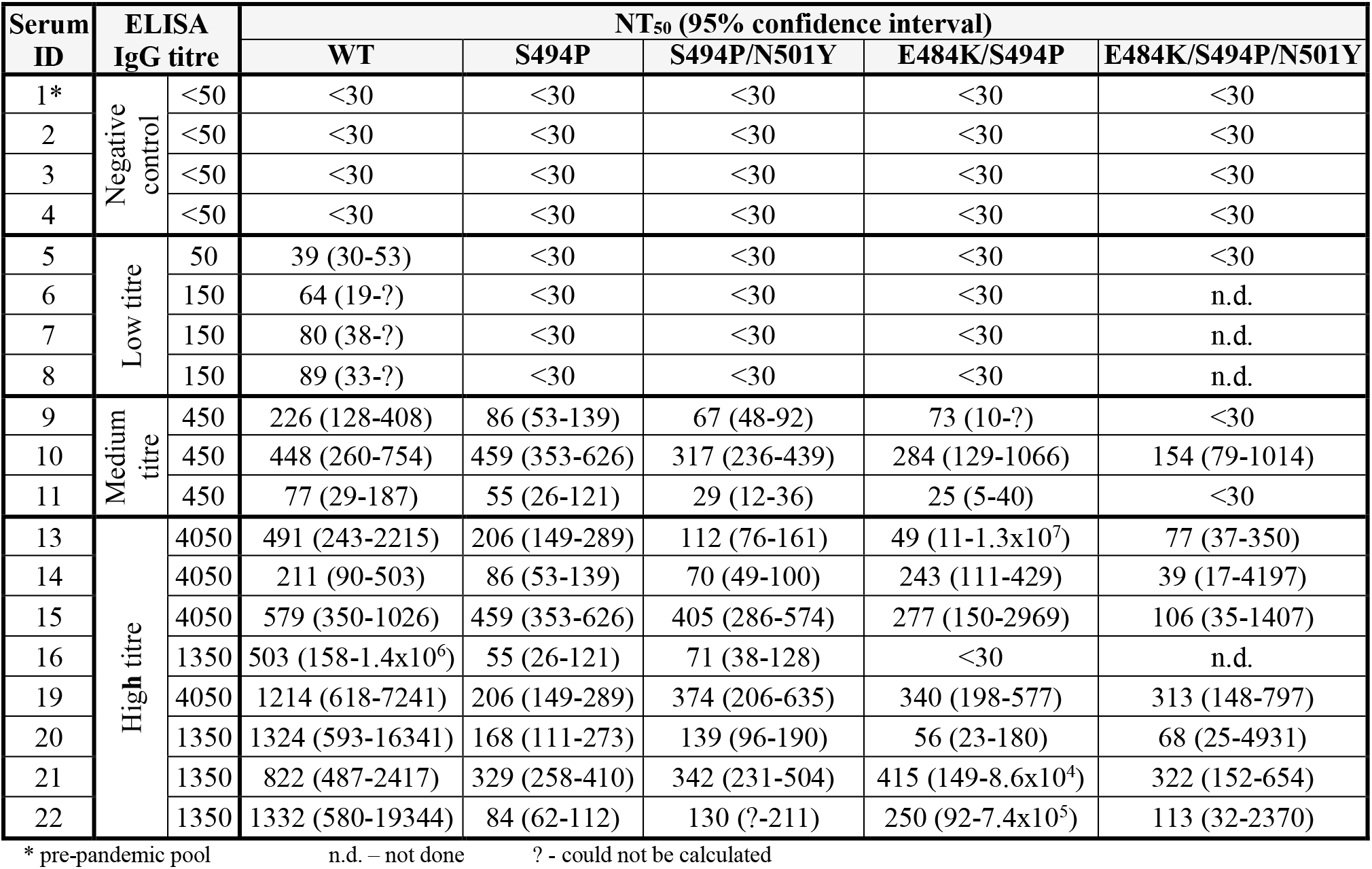
IgG antibody titres against SARS-CoV-2 spike protein and neutralizing titres (NT_50_) of convalescent sera against WT and mutant spike pseudoviruses.

**Supplementary Table 6.** IgG antibody titres against SARS-CoV-2 spike protein and neutralizing titres (NT_50_) against WT and mutant pseudoviruses of serum from vaccinated individuals, collected 1 month after the first and after the second doses of the vaccine.

| Serum ID | ELISA IgG titre | NT<sub>50</sub> (95% confidence interval) |  |  |  |  |
| --- | --- | --- | --- | --- | --- | --- |
|  |  | WT | S494P | S494P/N501Y | E484K/S494P | E484K/S494P/N501Y |
| A-1<sup>st</sup> | 150 | <30 | <30 | <30 | <30 | <30 |
| B-1<sup>st</sup> | 450 | <30 | <30 | <30 | <30 | <30 |
| H-1<sup>st</sup> | 36450 | 336 (116-?) | 278 (202-452) | 142 (99-229) | 284 (43-?) | 70 (25-?) |
| I-1<sup>st</sup> | 4050 | <30 | <30 | <30 | <30 | <30 |
| J-1<sup>st</sup> | 12150 | 60 (18-?) | 105 (56-1115) | 102 (50-1.5X10<sup>5</sup>) | 52 (16-105) | 44 (23-1364) |
| K-1<sup>st</sup> | 1350 | <30 | <30 | <30 | <30 | <30 |
| L-1<sup>st</sup> | 12150 | 219 (74-?) | 395 (307-505) | 332 (219-509) | 233 (42-?) | 74 (43-114) |
| M-1<sup>st</sup> | 36450 | 91 (51-819) | 37 (22-69) | 51 (22-19545) | 123 (36-?) | 49 (5-?) |
| H-2<sup>nd</sup> | 328050 | 631 (422-948) | 715 (?-909) | 495 (345-697) | 282 (143-575) | 210 (136-331) |
| I-2<sup>nd</sup> | 12150 | 263 (164-415) | 192 (?-251) | 122 (84-154) | 128 (57-746) | 43 (18-79) |
| J-2<sup>nd</sup> | 36450 | 191 (116-313) | 206 (?-292) | 154 (116-206) | 63 (19-1158) | 78 (39-221) |
| K-2<sup>nd</sup> | 36450 | 119 (70-213) | 140 (97-205) | 109 (74-151) | 69 (30-139) | 52 (0-?) |
| L-2<sup>nd</sup> | 36450 | 858 (556-1430) | 389 (300-517) | 221 (143-349) | 138 (57-462) | 118 (53-1860) |
| M-2<sup>nd</sup> | 36450 | 301 (122-2023) | 270 (202-353) | 264 (172-411) | 111 (59-220) | 170 (106-289) |
| N-2<sup>nd</sup> | 12150 | 97 (63-138) | 106 (70-164) | 77 (52-?) | 34 (21-49) | <30 |
| O-2<sup>nd</sup> | 36450 | 259 (125-670) | 142 (120-209) | 126 (96-162) | 77 (39-147) | 79 (38-146) |

? - could not be calculated

**Supplementary Table S7.**
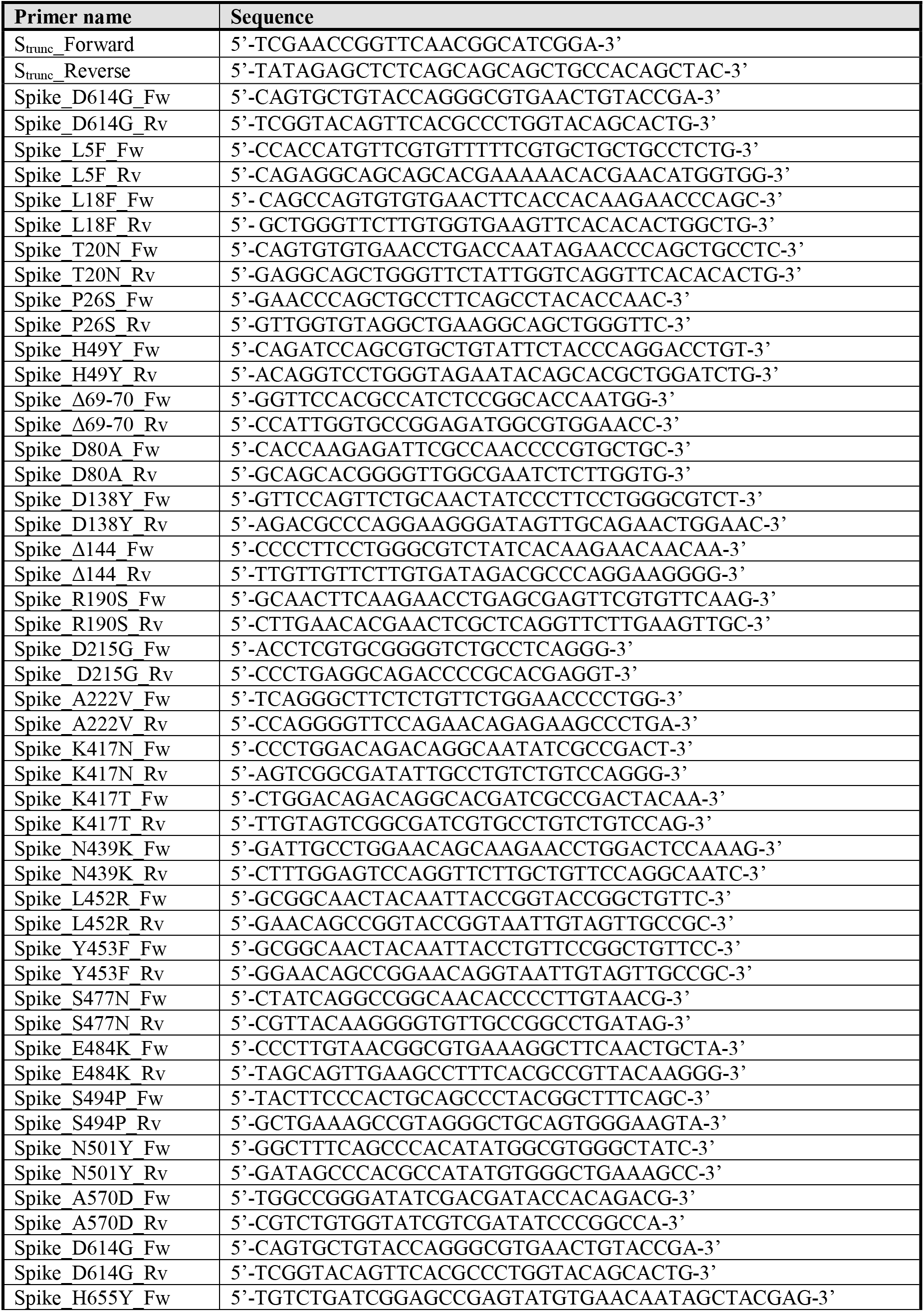
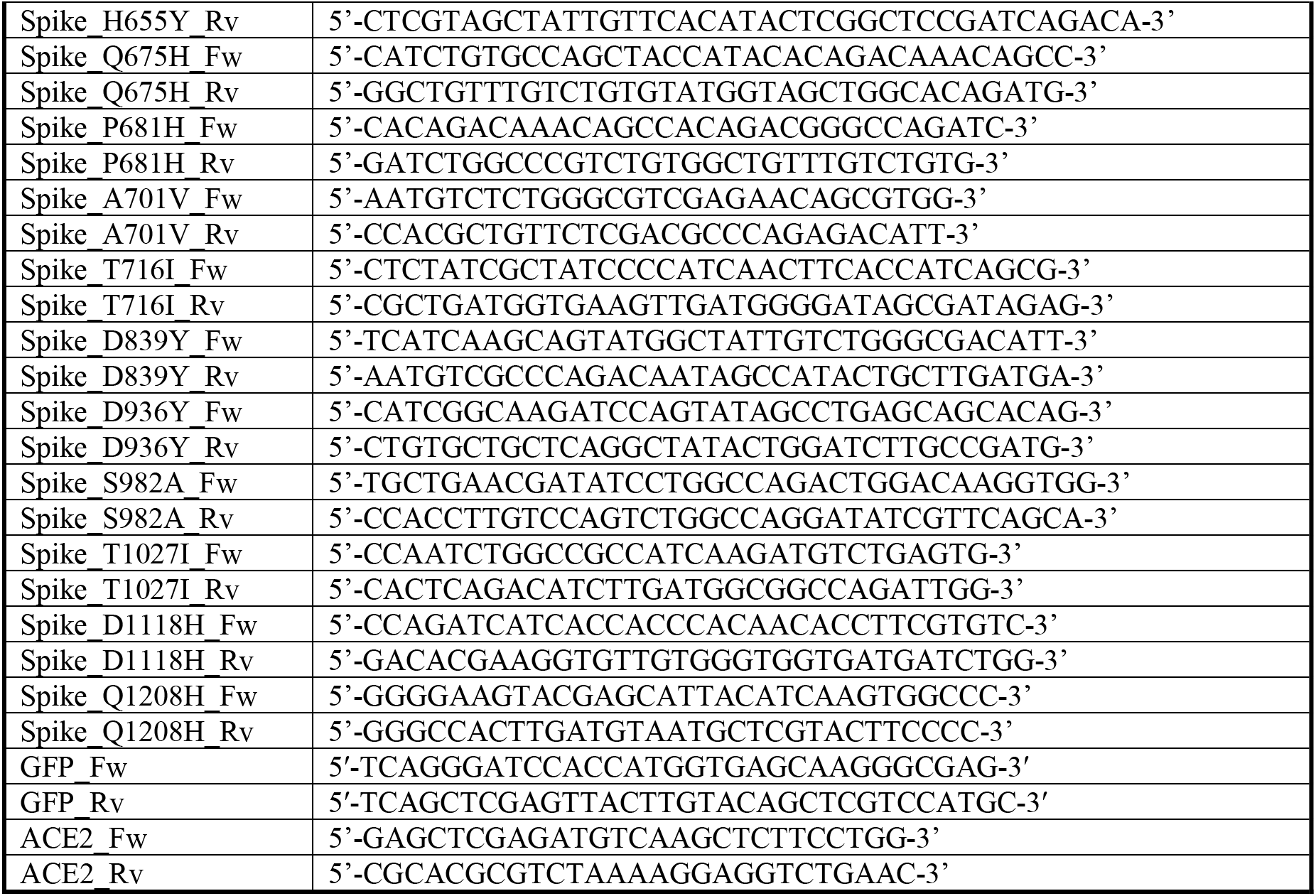
Primers used in this study.

